# Bacteria Hijack a Host Metabolite via Three Distinct Sensors to Orchestrate Infection

**DOI:** 10.64898/2026.06.19.733280

**Authors:** Qinmeng Liu, Shuyu Li, Yufei Zhao, Sidi Chen, Heyan Shen, Yayong Yang, Tao Huang, Yongdong Li, Zhiyan Wei, Zhuo Wang, Yi Chen, Lingfang Zhu, Yao Wang, Xihui Shen

## Abstract

The crisis of antimicrobial resistance demands new strategies that circumvent conventional killing. Here, we show that *Yersinia pseudotuberculosis* intercepts host mesifurane, a mimic of quorum-sensing signal AI-2. Rather than acting through a single receptor, mesifurane is directly and independently sensed by three receptor classes, YcrA, NarX, and CusR, that converge on a “Sense, Arm, Recruit” program. YcrA drives chemotaxis toward host cells; NarX-NarP licenses T6SS-mediated killing; and CusR amplifies AI-2 production to recruit more bacteria. YcrA is conserved across Gram-negative pathogens, creating a therapeutic vulnerability. Suramin sodium, a repurposed drug blocking YcrA, disarms this circuit without inhibiting growth. In murine sepsis, suramin sodium conferred 80–90% survival against WHO critical-priority pathogens, including carbapenem-resistant *A. baumannii*, *E. coli*, and *P. aeruginosa*, even when delayed 8 hours post-infection, without selecting for resistance. These findings establish that disrupting sensory perception, rather than viability, provides an anti-infective strategy bypassing resistance while imposing minimal selective pressure.

## Introduction

The escalating crisis of antimicrobial resistance has exposed the limits of conventional antibiotics. Few new classes of antibiotics have reached the clinic in recent decades, and for multidrug-resistant pathogens, which already account for over 1.2 million deaths annually^1,2^, treatment options are increasingly limited^3^.

Antivirulence strategies that disarm rather than kill bacteria have attracted sustained interest, yet their clinical impact has been modest^4,5^, in part because most efforts converge on a narrow set of recurring targets, such as quorum sensing, secretion systems, adhesins, toxins, and biofilms, each of which pathogens can circumvent through redundant pathways^6,7,8^. Overcoming this redundancy may require strategies that simultaneously disrupt multiple virulence programs.

Bacteria depend on environmental cues, including host-derived signals, to gauge their location within the host and activate appropriate virulence responses^9,10,11^. Interfering with the perception of such signals could, in principle, disable multiple downstream programs at once^6,12,13^. Yet few host-derived signals capable of orchestrating a coordinated virulence program have been identified, and whether any single host signal can trigger a multi-module offensive program has remained unknown.

A critical setting where such signals likely operate is the encounter between bacteria and immune cells. Macrophages patrol infected tissues and destroy invaders^14^, but some pathogens have turned the tables. *Vibrio cholerae*, for example, assembles biofilm-like structures on human phagocytes to destroy them as a multicellular predator^15^. While most pathogens evade immune cells through more indirect means^16,17,18^, the ability to sense these cells is probably widespread. This raises a fundamental question: how does a bacterium recognize that it is confronting an immune cell, and what cues initiate the response?

One intriguing class of host-derived signals comprises structural mimics of bacterial autoinducer-2 (AI-2), a quorum-sensing molecule that mediates interspecies communication^19,20,21^. Such mimics have been found in yeast^22^, plants^23^, and mammals^24,25^, and some have been chemically characterized. However, whether and how these host-derived mimics function during infection has remained unclear. They could interfere with bacterial signaling or be co-opted by pathogens as cues to activate virulence. Distinguishing between these possibilities requires identifying the host-derived factor and determining how bacteria perceive and respond to it.

Here we show that the enteric pathogen *Yersinia pseudotuberculosis* (*Yptb*) co- opts host AI-2 mimics for offense. We identify mesifurane as a mammalian AI-2 mimic that triggers a multisensory circuit through which the bacterium executes a coordinated “Sense–Arm–Recruit” response, involving chemotaxis (YcrA), T6SS licensing (NarX–NarP), and population recruitment (CusR). This circuit is required for colonization and virulence. We further show that suramin sodium, an approved drug^26,27^, disrupts this circuit by antagonizing YcrA and protects mice from lethal infection by carbapenem-resistant *A. baumannii*, *E. coli*, and *P. aeruginosa*, all WHO critical-priority pathogens^28^, even when treatment was delayed until 8 hours after infection. These findings establish that intercepting pathogen sensory perception, rather than viability, defines an anti-infective strategy that bypasses conventional resistance and offers a path forward against multidrug-resistant pathogens.

## Result

### A host-derived AI-2 mimic guides bacterial chemotaxis toward macrophages

The initial encounter between the pathogen and host often dictates the outcome of infection. We found that wild-type (WT) *Yptb* exhibited a striking tropism for macrophages, forming dense, biofilm-like aggregates on the surface of RAW 264.7 murine macrophages (Figures 1A and 1B), reminiscent of the invasive biofilms recently reported for *Vibrio cholerae* on macrophages^15^. This interaction was far from passive: in Transwell assays, *Yptb* migrated robustly and directionally toward the macrophages, indicating active chemotaxis (Figures 1C and 1D). Time-lapse tracking of GFP-labeled bacteria confirmed their preferential accumulation in macrophage- containing chambers, revealing a clear chemoattractant gradient (Figure S1A).

**Figure 1.**
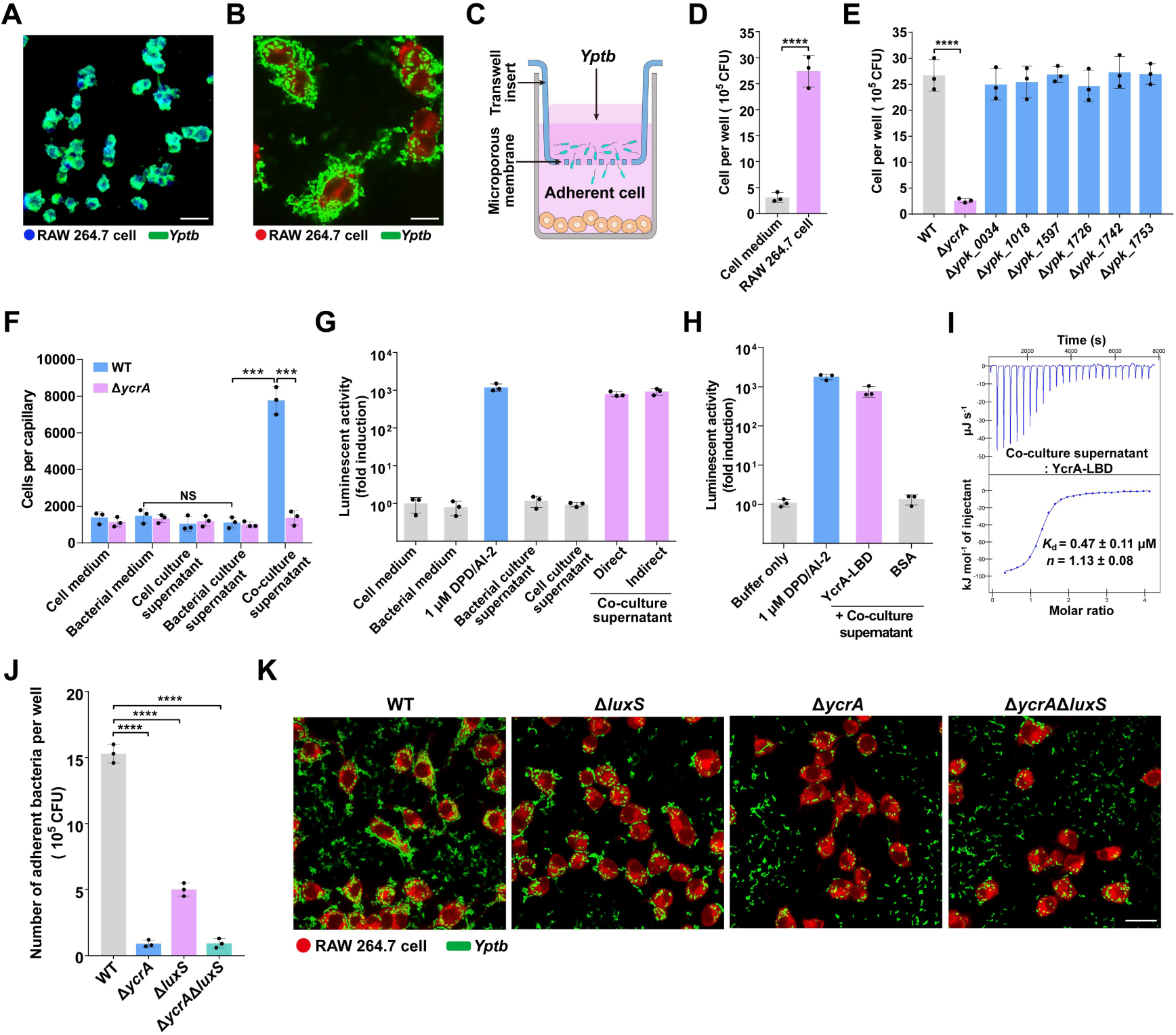
Macrophage-derived AI-2 mimic is sensed by YcrA to drive *Yptb* chemotaxis and adhesion. **(A, B)** Microscopic visualization of *Yptb* adhesion to macrophages. Adhesion of GFP- expressing *Yptb* to RAW 264.7 macrophages (MOI 100) was visualized by inverted fluorescence microscopy (A) and confocal microscopy (B). GFP-expressing *Yptb* (green) and macrophages were stained with DAPI (blue; A) or CellTracker™ Deep Red (red; B). Scale bars: 100 μm (A) and 20 μm (B). Images are representative of three independent experiments with similar results. **(C)** Schematic of the transwell chemotaxis assay. A bacterial suspension was added to the upper chamber, separated by a porous membrane (5 μm) from a macrophage monolayer in the lower chamber. **(D)** Quantification of WT *Yptb* migration toward RAW 264.7 macrophages in the transwell assay. Bacteria that translocated to the lower chamber after 2 h were enumerated by CFU plating. Sterile cell medium (cell-free) served as a control. Data are shown as CFU counts. **(E)** Transwell migration of WT and indicated chemoreceptor mutant strains toward macrophages. Migrated bacteria were quantified, and final CFU counts were corrected by subtracting the number of bacteria that migrated into cell-free control wells. **(F)** Chemotactic response of WT and Δ*ycrA Yptb* to cell medium, bacterial medium, cell culture supernatant, bacterial culture supernatant, or host-bacterial co-culture supernatant, assessed by quantitative capillary assay. Co-culture supernatant was prepared with Δ*luxS Yptb* to eliminate bacterially produced AI-2, thus ensuring any activity originates from the host. The data represent bacterial accumulation in capillaries. **(G)** AI-2 mimic activity is detected only upon co-culture of macrophages with *Yptb*. RAW 264.7 macrophages were stimulated with *Yptb* Δ*luxS* either by direct contact or via non-contact (separated by a semipermeable membrane). After 5 h, AI-2 mimic activity in filter-sterilized supernatants was measured using the *Vibrio harveyi* MM32 bioluminescence reporter assay. Cell medium, bacterial medium, and supernatants from *Yptb* Δ*luxS* cultured alone or RAW 264.7 cells cultured alone showed no detectable activity. DPD/AI-2 (1 μM) served as a positive control. Data are presented as fold induction relative to buffer control. **(H)** Ligand capture assay showing binding of the AI-2 mimic to YcrA. Purified YcrA- LBD was incubated with co-culture supernatant; the captured ligand was heat-eluted and quantified using the *V. harveyi* MM32 reporter assay. BSA served as a negative control. Data are fold induction relative to the buffer control. **(I)** AI-2 mimic binds YcrA with high affinity. ITC assay was conducted to determine the specific interaction between YcrA-LBD and the co-culture supernatant. The displayed thermogram is one representative of three independent experiments; *K*^d^ and binding stoichiometry (*n*) are presented as mean ± s.d. from three independent experiments. **(J)** RAW 264.7 macrophages were infected with the indicated *Yptb* strains at an MOI of 100 for 1 h. Nonadherent bacteria were removed by washing, and adherent bacteria were released by cell lysis, serially diluted, and enumerated by CFU plating. **(K)** RAW 264.7 macrophages (CellTracker™ Deep Red, red) were infected with GFP-expressing *Yptb* strains (green) at MOI 100 for 1 h. Nonadherent bacteria were removed by washing prior to imaging. Scale bar, 20 μm. Images are representative of three independent experiments. Data are represented as mean ± SD of three biological replicates, each with three technical replicates. Statistical significance was determined with the two-tailed unpaired Student’s *t*-test. \*\*\**P* < 0.001; \*\*\*\**P* < 0.0001; NS, not significant.

To identify the bacterial receptor mediating this host-directed navigation, we systematically screened *Yptb* mutants lacking each of its seven methyl-accepting chemotaxis proteins (MCPs)^29^. Deletion of a single gene, *ypk_2833*, which we named *ycrA*, abolished macrophage-directed migration, reducing it by more than 90% compared with the WT control (Figure 1E). By contrast, loss of any of the other six MCPs had no measurable impact on chemotaxis. This chemotactic defect was fully rescued by plasmid-based complementation, establishing YcrA as the cognate receptor for this behavior (Figure S1B). Importantly, the growth kinetics of the Δ*ycrA* mutant were indistinguishable from those of WT *Yptb* (Figure S1C), ruling out impaired fitness as the cause of the observed migration defect.

Given that YcrA was required for macrophage-directed chemotaxis, we reasoned that its cognate ligand should be present in the macrophage environment. However, supernatant from macrophage monocultures failed to attract *Yptb* (Figure S1D), suggesting that the relevant signal is not constitutively secreted by macrophages alone. We therefore tested cocultures of macrophages with WT *Yptb*. This coculture supernatant strongly stimulated bacterial migration (Figure S1D). Notably, supernatant from *Yptb* cultured alone also exhibited weak chemotactic activity (Figure S1D). Because AI-2 produced by many enteric bacteria is itself a known chemoattractant^21,30,31^, we suspected that the residual activity in the bacterial monoculture supernatant might reflect endogenously produced AI-2. To eliminate this confounding variable, we turned to a *Yptb* Δ*luxS* mutant, which cannot produce AI- 2^30,32^. Supernatant from the Δ*luxS* mutant cultured alone completely lost its chemoattractant activity (Figure 1F), confirming that the weak attraction seen with WT bacterial supernatant was indeed attributable to AI-2. Crucially, supernatant harvested from cocultures of macrophages with the Δ*luxS* mutant retained potent chemotactic activity (Figure 1F). This activity was entirely dependent on YcrA, as it was absent in the Δ*ycrA* strain and was rescued upon complementation (Figures 1F and S1E). These results suggest that bacterial interaction induces the production or release of a host-derived factor that acts through YcrA.

To investigate how YcrA recognizes the host-derived signal, we next characterized its ligand-binding properties. Bioinformatic analysis revealed that YcrA harbors a dCache-type ligand-binding domain (LBD), a hallmark of chemoreceptors that sense small molecules such as organic acids and AI-2^21,33,34^ (Figure S1F). Sequence alignment revealed that YcrA shares critical conserved residues with known AI-2 receptors^21,34^ (Figure S1G). Biochemical experiments confirmed this prediction: compared with a control protein, recombinant YcrA-LBD selectively increased AI-2 activity >1,000-fold, and isothermal titration calorimetry (ITC) revealed a high-affinity interaction between YcrA and AI-2 (Figures S1H and S1I). *Yptb* also displayed YcrA-dependent chemotaxis toward AI-2 (Figure S1J).

We therefore hypothesized that the macrophage-derived attractant might be a structural mimic of AI-2^24^. Indeed, the coculture supernatant robustly activated a canonical *Vibrio harveyi* AI-2 reporter strain^21,24^ (Figure 1G). This activity was recapitulated in transwell setups where bacteria and macrophages were physically separated, indicating that the soluble factor, rather than contact-dependent signaling, sufficed for this effect (Figure 1G). Cell-free bacterial culture supernatant, even when added at a volume as low as 10 µL to a 2 mL culture of macrophages (a 200-fold dilution), was sufficient to stimulate AI-2 mimic production (Figure S1K), ruling out nonspecific effects from bacterial growth and directly implicating a secreted bacterial factor that host cells sense at exceedingly low levels. Ligand capture assays demonstrated specific binding of this mimic to YcrA (Figure 1H), and ITC revealed a direct, high-affinity interaction (Figure 1I). In contrast, binding of the supernatant from macrophages cultured alone was not detectable (Figure S1L), confirming that the signal is host-derived but dependent on bacterial stimulation.

We next examined the functional consequences of this YcrA-mediated sensing. Deletion of *ycrA* severely impaired the stable adhesion of *Yptb* to macrophages (Figure 1J). The loss of endogenous AI-2 production (Δ*luxS*) moderately reduced adhesion, yet the Δ*ycrA*Δ*luxS* double mutant phenocopied the Δ*ycrA* single mutant (Figure 1J), indicating that YcrA was the dominant pathway. High-resolution confocal microscopy further confirmed that AI-2 mimic-induced bacterial clustering and adhesion on macrophage surfaces were strictly YcrA dependent (Figure 1K). Thus, a host-produced AI-2 mimic serves as the primary chemoattractant, guiding *Yptb* to its target via YcrA, with endogenous bacterial AI-2 playing a secondary, reinforcing role in adhesion. The identity of this host-derived signal, and whether it represents a known metabolite or a novel AI-2 analog, remained to be determined.

### Mesifurane was identified as a conserved mammalian AI-2 mimic

Having established that a host-derived AI-2 mimic acts through YcrA (Figure 1), we next sought to identify this chemoattractant at the molecular level using a multi-pronged strategy that combined metabolomic profiling with biochemical analyses. To minimize confounding effects from bacterial metabolites, we first stimulated macrophages with phosphate-buffered saline (PBS), a treatment previously reported to elicit AI-2 mimic activity^24^. PBS-treated RAW 264.7 cells indeed exhibited such activity (Figure S2A), promoting us to perform untargeted liquid chromatography‒tandem mass spectrometry (LC‒MS/MS) metabolomic profiling on their supernatants. This analysis detected 943 metabolic features in PBS-treated macrophage supernatant and 1,256 in those from cocultures with *Yptb ΔluxS* (Table S1), providing the foundational dataset for subsequent identification of the host-derived AI-2 mimic.

To identify potential AI-2 mimics among these metabolic features, we developed a bioinformatic pipeline. We annotated overlapping features against the Human Metabolome Database and evaluated their binding to YcrA-LBD using deep learning- guided molecular docking. Candidates with high predicted binding affinity (docking score ≤ −7.0 kcal mol⁻^1^) were prioritized (Table S2). Given the confirmed affinity of YcrA for both AI-2 and the mimic, we restricted our analyses to metabolites with structural and mass similarity to DPD/AI-2. Among these compounds, only one— mesifurane (2,5-dimethyl-4-methoxy-2,3-dihydro-3-furanone)—potently activated the AI-2 reporter strain (Figure 2A).

**Figure 2.**
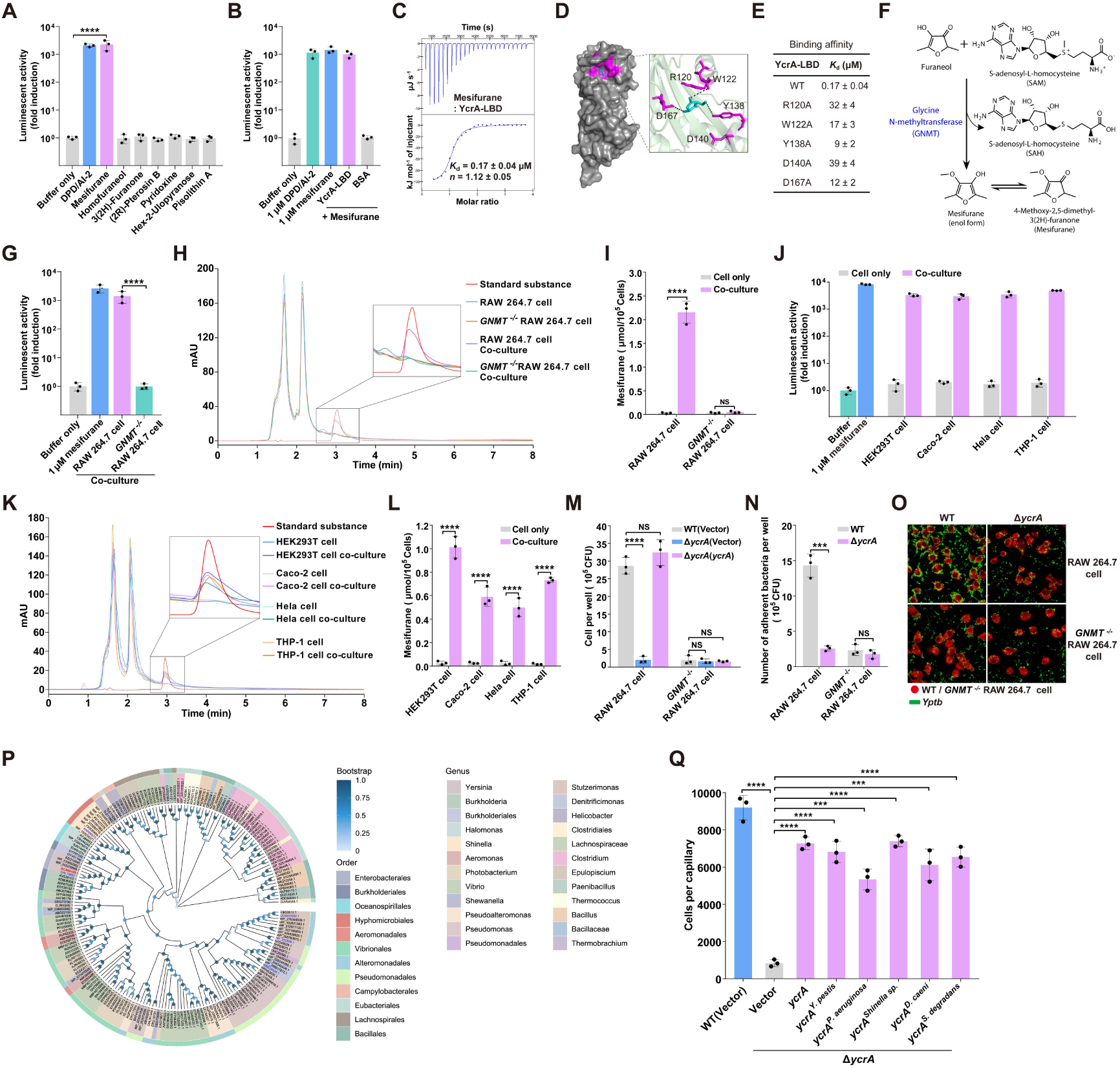
Mesifurane is the mammalian AI-2 mimic recognized by YcrA. **(A)** Mesifurane stimulates light production in *V. harveyi* MM32 reporter strain. Bioluminescence in MM32 by 1 μM mesifurane, Homofuraneol, 3(2H)-Furanone, (2R)-Pterosin B, Pyridoxine, Hex-2-Ulopyranose, and Pisolithin A. 1 μM DPD/AI-2 served as a positive control. Data are shown as fold induction relative to a buffer control. **(B)** YcrA specifically captures mesifurane. Purified YcrA-LBD was incubated with mesifurane; the captured ligand was heat-eluted and quantified using the *V. harveyi* MM32 reporter assay. BSA was used as a negative control. Data are fold induction relative to the buffer control. **(C)** ITC analysis of the specific interaction between YcrA-LBD and mesifurane. The displayed thermogram is one representative of three independent experiments; *K*^d^ and binding stoichiometry (*n*) are presented as mean ± s.d. from three independent experiments. **(D)** Predicted binding mode of mesifurane within the YcrA-LBD. The optimal docking pose of mesifurane (cyan sticks) was determined using AutoDock4.42 and Chimera. Key interacting residues are shown as purple sticks, with dashed lines indicating potential hydrogen bonds. **(E)** Binding of mesifurane to YcrA-LBD and its mutants. The binding affinity was measured by ITC. The *K*^d^ values are presented as mean ± s.d. of three independent experiments. **(F)** Schematic representation of mesifurane biosynthesis via GNMT (glycine N- methyltransferase). SAM serves as the methyl donor for the methylation of furaneol by GNMT, yielding mesifurane and S-adenosylhomocysteine (SAH). Mesifurane and its enol form interconvert via keto–enol tautomerism. **(G)** *GNMT*^-/-^ RAW 264.7 cells fail to produce mesifurane. Δ*luxS Yptb* was co-cultured with WT or *GNMT*^-/-^ RAW 264.7 cells, and mesifurane activity was measured using the *V. harveyi* MM32 reporter. 1 μM mesifurane as a positive control. Data are fold induction relative to the buffer control. **(H)** HPLC analysis of mesifurane in supernatants. Overlaid chromatograms show the mesifurane standard (red), co-culture supernatants from WT (purple) and *GNMT*^-/-^ (green) RAW 264.7 cells, and cell culture supernatants from WT (blue) and *GNMT*^-/-^ (orange) RAW 264.7 cells. The active fraction is enlarged (inset). mAU, milli- absorbance units. **(I)** Quantification of mesifurane in supernatants by LC-MS/MS. Mesifurane levels were measured from WT and *GNMT*^-/-^ RAW 264.7 cells under co-culture and non-co- culture conditions. **(J)** Multiple cell lines can all be stimulated to produce mesifurane. Supernatants from co-cultures of Δ*luxS Yptb* with the indicated human cell lines (HEK293T, Caco-2, HeLa, THP-1) were assayed for mesifurane activity using the *V. harveyi* MM32 reporter. 1 μM mesifurane was a positive control. Supernatants from mammalian cells cultured alone served as negative controls. Data are fold induction relative to the buffer control. **(K)** HPLC profiling of mesifurane in various mammalian cell supernatants. Overlaid chromatograms show the mesifurane standard (red) and co-culture (dark hues) versus cell culture (light hues) supernatants from the indicated cell lines, with the active fraction enlarged (inset). mAU, milli-absorbance units. **(L)** LC-MS/MS quantification of mesifurane from different mammalian cell lines. Mesifurane levels were measured under co-culture and non-co-culture conditions for each cell type. **(M)** Chemotaxis of *Yptb* toward macrophages requires host mesifurane production. Transwell assays quantifying migration of the indicated strains toward WT or *GNMT*_-/-_ RAW 264.7 cells. Migrated bacteria were quantified, and final CFU counts were corrected by subtracting the number of bacteria that migrated into cell-free control wells. **(N)** WT and *GNMT*_-/-_ RAW 264.7 cells were infected with the indicated *Yptb* strains at MOI 100 for 1 h. Nonadherent bacteria were removed by washing, and adherent bacteria were released by cell lysis, serially diluted, and enumerated by CFU plating. **(O)** WT and *GNMT*_-/-_ RAW 264.7 cells (CellTracker™ Deep Red, red) were infected with GFP-expressing *Yptb* strains (green) at MOI 100 for 1 h. Nonadherent bacteria were removed by washing prior to imaging. Scale bar, 20 μm. Images are representative of three independent experiments. **(P)** Phylogenetic analysis of YcrA homologs. A maximum-likelihood tree was constructed from 200 YcrA sequences. The *Yptb* YcrA is highlighted in red, and experimentally verified homologs are in blue. Outer rings indicate bacterial taxonomy at the order level. Bootstrap values are visualized as a blue gradient. **(Q)** Heterologous complementation with *ycrA* orthologs restores mesifurane chemotaxis in *Yptb* Δ*ycrA*. Chemotaxis of the indicated strains toward 10 μM mesifurane was assessed by quantitative capillary assay. *Yptb* Δ*ycrA* was complemented with *ycrA* homologous genes from *Yersinia pestis* CO92, *Pseudomonas aeruginosa* PAO1, *Shinella sp.* HZN7, *Denitrificimonas caeni,* and *Stutzerimonas degradans*. Accumulated cells were quantified and corrected by subtracting background migration into PBS-containing capillaries. Data are represented as mean ± SD of three biological replicates, each with three technical replicates. Statistical significance was determined with the two-tailed unpaired Student’s *t*-test. \*\*\**P* < 0.001; \*\*\*\**P* < 0.0001; NS, not significant.

We next sought to validate mesifurane as the bona fide ligand for YcrA using complementary biochemical and functional assays. In ligand-release assays, the YcrA-LBD bound to mesifurane and subsequently activated the AI-2 reporter with a potency equivalent to that of 1 µM DPD/AI-2 (Figure 2B). ITC measurements confirmed a direct, high-affinity interaction between YcrA and mesifurane (*K*^d^ *=* 0.17 ± 0.04 μM) (Figure 2C). Molecular docking simulations revealed mesifurane within the YcrA binding pocket, forming close contacts with YcrA-LBD residues R120, W122, Y138, D140, and D167 (Figure 2D). Alanine substitution at these positions dramatically reduced binding affinity, confirming their critical role (Figures 2E and S2B). Functionally, purified mesifurane elicited robust, YcrA-dependent chemotaxis (Figure S2C). These results establish mesifurane as a bona fide ligand for YcrA.

Since YcrA recognizes both AI-2 and mesifurane, we investigated how bacteria respond when both signals are present simultaneously—as they would be during infection. We first determined the physiological concentration of each signal. WT *Yptb* produced AI-2 in a growth phase-dependent manner, peaking at ∼6 h and reaching 5–10 μM in culture supernatants (Figures S2D and S2E). Mesifurane reached similar levels (5–10 μM) in macrophage coculture supernatants (Figure S2F). We therefore compared chemotactic responses across this shared concentration range (1–10 μM). Compared with AI-2, mesifurane elicited stronger chemotaxis at every matched concentration, and the response to 1 μM mesifurane was comparable to that evoked by 5 μM AI-2 (Figure S2G). This hierarchical preference ensures that *Yptb* prioritizes the host-derived signal even when its own quorum-sensing autoinducer is abundant.

Having established mesifurane as the dominant host-derived ligand, we next investigated its biosynthetic origin. Metabolic database mining suggested that mesifurane could arise as a product of a specific S-adenosylmethionine (SAM)- dependent methyltransferase, GNMT (glycine N-methyltransferase; KEGG ID: 14711) reaction, generating S-adenosylhomocysteine (SAH) and mesifurane (Figure 2F). Using CRISPR-Cas9^35^, we generated a knockout of GNMT in RAW 264.7 cells (*GNMT^-/-^*) (Figure S2H and Table S3). These mutant cells completely lost the ability to produce mesifurane during coculture (Figure 2G). Subsequent HPLC and targeted LC‒MS/MS analyses confirmed that mesifurane was detectable exclusively in supernatants from cocultures involving WT macrophages, with no signal from *GNMT^-/-^* cocultures or any macrophage monoculture (Figures 2H, 2I and S2I). Thus, host- derived mesifurane is produced via a GNMT‒dependent biosynthetic pathway in response to bacterial encounter.

This response was not restricted to macrophages. Multiple human and murine cell lines—including monocytic THP-1, epithelial Caco-2, HEK293T, and HeLa cells—produced detectable mesifurane upon coculture with *Yptb* Δ*luxS* (Figures 2J- 2L and S2J). Moreover, the trigger was not unique to *Yptb*; diverse bacterial pathogens, including *Pseudomonas aeruginosa*, *Vibrio harveyi*, *Salmonella* Typhimurium, *Escherichia coli*, and *Klebsiella pneumoniae*, all induced mesifurane production in host cells (Figures S2K-S2N). This finding identifies mesifurane as a conserved host-derived chemoattractant, a metabolite released in response to bacterial interaction that serves as a cue for bacterial virulence.

We next asked whether host-derived mesifurane was required for *Yptb* navigation and adhesion. In transwell assays, *GNMT^-/-^* macrophages failed to attract *Yptb* (Figure 2M). Accordingly, bacterial adhesion was significantly reduced on *GNMT^-/-^* compared with WT macrophages, whereas the adhesion of the Δ*ycrA* mutant was uniformly low on both (Figures 2N and 2O). These results establish host-derived mesifurane as an essential cue for YcrA-dependent navigation and adhesion.

Phylogenetic analysis revealed that YcrA orthologs carrying the key mesifurane- binding residues are widely distributed across diverse bacterial genera, including *Yersinia pestis*, *Pseudomonas aeruginosa*, *Shinella zoogloeoides*, *Denitrificimonas caeni*, and *Stutzerimonas degradans* (Figures 2P and S2O). Notably, heterologous expression of the corresponding *ycrA* genes from these species in the *Yptb* Δ*ycrA* mutant fully restored mesifurane sensing and chemotaxis (Figure 2Q). These findings indicate that the ability to recognize this host-derived signal via YcrA is a widely distributed bacterial trait. Collectively, these results establish mesifurane as a conserved mammalian signal that is broadly recognized by bacterial YcrA orthologs.

### Mesifurane transcriptionally reprograms bacterial virulence and activates T6SS-mediated cytotoxicity

Having established that mesifurane guides *Yptb* to macrophages, we next asked whether this host metabolite does more than attract, specifically, whether it also transcriptionally reprograms bacterial virulence upon arrival. RNA sequencing revealed that bacterial exposure to mesifurane triggered extensive transcriptional remodeling, with 736 genes upregulated and 84 downregulated (|log^2^FC| ≥ 1, FDR < 0.05) (Figure 3A). Pathway analysis revealed strong enrichment for processes essential for host engagement, such as chemotaxis, bacterial secretion systems, two-component signaling, flagellar assembly, quorum sensing, and biofilm formation (Figure 3B). Within the secretion systems, the type VI secretion system (T6SS) drew our particular attention. Strikingly, all core genes of the T6SS-4 cluster were significantly induced (Figure 3C). qRT‒PCR confirmed that six core T6SS-4 genes, along with biofilm- and flagellum-associated genes, were significantly upregulated by either the coculture supernatant or purified mesifurane (Figures 3D and S3A-S3C). These latter genes are integral to bacterial motility and surface adhesion, suggesting that mesifurane coordinates multiple programs essential for host engagement, from motility and attachment to the deployment of the T6SS.

**Figure 3.**
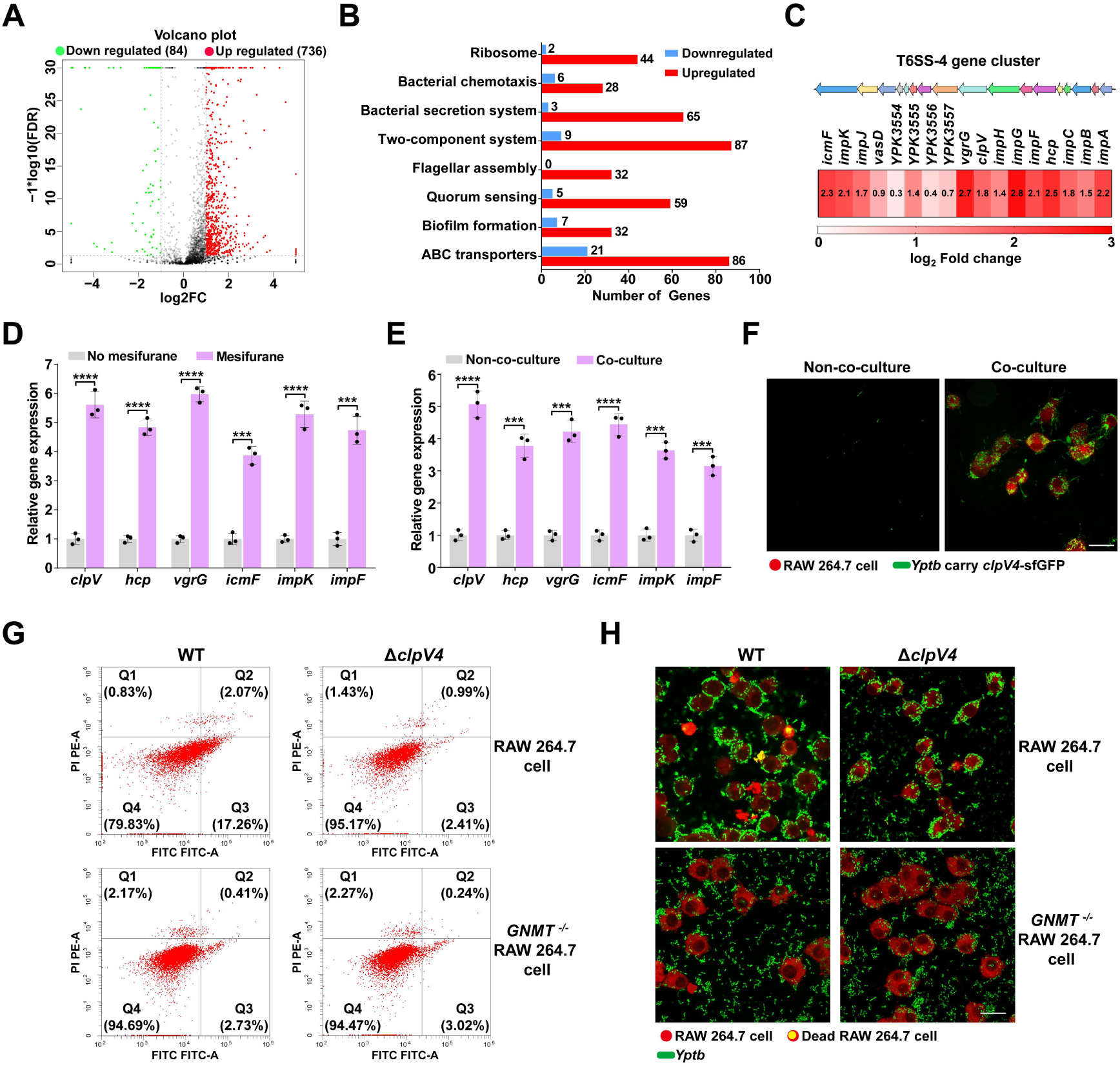
Mesifurane induces *Yptb* T6SS-4 and enhances macrophage cytotoxicity. **(A)** Transcriptomic analysis of *Yptb* in response to mesifurane. Volcano plot of RNA- seq data from *Yptb* incubated with host-bacteria co-culture supernatant versus untreated control for 2 h. Differentially expressed genes (DEGs) were identified based on two criteria (|fold change| ≥ 2 and false discovery rate (FDR) < 0.05). The x-axis shows the log2 (fold change) in gene expression, while the y-axis depicts the negative log10 of the FDR. The black dashed vertical lines indicate the 2-fold change thresholds, expressed as [log2 (-1, +1)]. The black dashed horizontal line represents the FDR = 0.05, expressed as [-log10 (1.30)]. **(B)** Gene Ontology (GO) enrichment analysis of DEGs. The top eight significantly enriched terms in the Biological Process, Cellular Component, and Molecular Function categories are shown. Enrichment was computed separately for upregulated (red) and downregulated (blue) DEGs (|fold change| ≥ 2). **(C)** Transcriptional activation of the T6SS-4 gene cluster by the mesifurane. Gene organization of the T6SS-4 cluster and heat map showing the expression of all differentially transcribed genes in the T6SS-4 cluster in WT *Yptb* treated with co- culture supernatant versus untreated control, as determined by RNA-seq. **(D)** qRT-PCR analysis of T6SS core gene expression in *Yptb* left untreated or stimulated with 10 μM mesifurane for 2 h. Expression was normalized to 16S rRNA and is presented as fold change relative to the untreated control. **(E)** T6SS gene induction during macrophage infection. qRT-PCR analysis of T6SS gene expression in WT *Yptb* following infection of RAW 264.7 macrophages. Bacteria were added to macrophages at an MOI of 100, and after 2 h of co-culture, bacterial RNA was extracted for analysis. Expression was normalized to 16S rRNA and is presented as fold change relative to bacteria cultured without macrophages. **(F)** T6SS activity is induced in *Yptb* upon macrophage co-culture. Confocal micrographs of a reporter *Yptb* strain expressing a chromosomal ClpV4-sfGFP fusion (green). ClpV4 is the T6SS sheath-dismantling ATPase, and its focal accumulation indicates sites of active T6SS assembly. Bacteria were cultured alone (left) or co- cultured with RAW 264.7 macrophages (right, CellTracker™ Deep Red, red) at MOI 100 for 2 h. Scale bar, 20 μm. Images are representative of three independent experiments. **(G)** Flow cytometric analysis of macrophage apoptosis. WT and *GNMT*^-/-^ RAW 264.7 cells were infected with WT or Δ*clpV4 Yptb* (MOI 100) for 2 h, then stained with Annexin V-FITC and PI. A representative result from three independent experiments is shown. **(H)** Confocal microscopy assessment of macrophage death. Confocal micrographs of WT and *GNMT*^-/-^ RAW 264.7 cells (CellTracker™ Deep Red, red) were infected with GFP-tagged *Yptb* strains (green) at MOI 100 for 2 h. Dead macrophages were stained with PI (yellow). Scale bar, 20 μm. Images are representative of three independent experiments with similar results. Data are represented as mean ± SD of three biological replicates, each with three technical replicates. Statistical significance was determined with the two-tailed unpaired Student’s *t*-test. \*\*\**P* < 0.001; \*\*\*\**P* < 0.0001.

We next asked whether endogenous bacterial AI-2 could similarly activate the T6SS. In a Δ*luxS* mutant, supplementing AI-2 at a concentration exceeding the *Yptb* physiological peak (10 μM) (Figures S2D and S2E) failed to induce T6SS-4 genes (Figure S3D). In contrast, the host-derived mimic, mesifurane, triggered robust upregulation at the identical concentration (Figure S3D). Thus, mesifurane functions as a selective cue that coordinates motility, adhesion, and attack programs. Notably, the bacterial T6SS is licensed exclusively by the host-derived signal, not by the bacterial quorum-sensing molecule at physiologically relevant levels.

Given that the T6SS is a contact-dependent apparatus that can induce host cell apoptosis^36,37,38^, we hypothesized that YcrA-driven chemotaxis delivers bacteria to the host cell surface, where T6SS-4 is deployed to mediate killing. Consistent with this hypothesis, T6SS-4 gene expression was strongly induced during macrophage infection (Figure 3E). Using a fluorescent reporter for T6SS assembly (ClpV4- sfGFP)^39^, we observed bright, punctate fluorescence specifically at bacterium‒host contact sites (Figure 3F), confirming that host cell proximity triggers immediate structural assembly of the bacterial T6SS.

We next tested whether assembly of the bacterial T6SS leads to host cell death. Infection with WT *Yptb* induced substantial macrophage apoptosis (∼19.3%), whereas a T6SS-deficient mutant (Δ*clpV4*) caused minimal apoptosis (∼3.4%) (Figure 3G). Crucially, apoptosis was also abolished when WT *Yptb*-infected *GNMT^-/-^* macrophages, which cannot produce mesifurane, providing genetic evidence that host metabolite sensing is required for T6SS-mediated killing (Figure 3G). Similarly, propidium iodide (PI) staining demonstrated that macrophage death required both a functional T6SS and a host-derived mesifurane (Figure 3H). Thus, mesifurane not only draws the pathogen to its target but also licenses the lethal weaponry that eliminates the host cell. Host-derived mesifurane therefore functions as a dual- purpose signal, coupling bacterial navigation directly to virulence execution.

### The NarX–NarP two-component system directly senses mesifurane to activate the T6SS

To understand how the mesifurane signal is transduced to activate the T6SS, we performed a systematic genetic screen. We screened a library of *Yptb* mutants, each lacking one of its 25 sensor histidine kinases^39^, for their ability to activate a *P^T6SS4::^lacZ* reporter in response to mesifurane. Only the deletion of *narX* (*ypk_2232*) abolished this induction (Figure 4A). Accordingly, the mesifurane-induced expression of T6SS-4 core genes was completely eliminated in Δ*narX* (Figures 4B and S4A). Promoter-reporter assays confirmed that Δ*narX* failed to respond, and complementation restored inducibility (Figure S4B).

**Figure 4.**
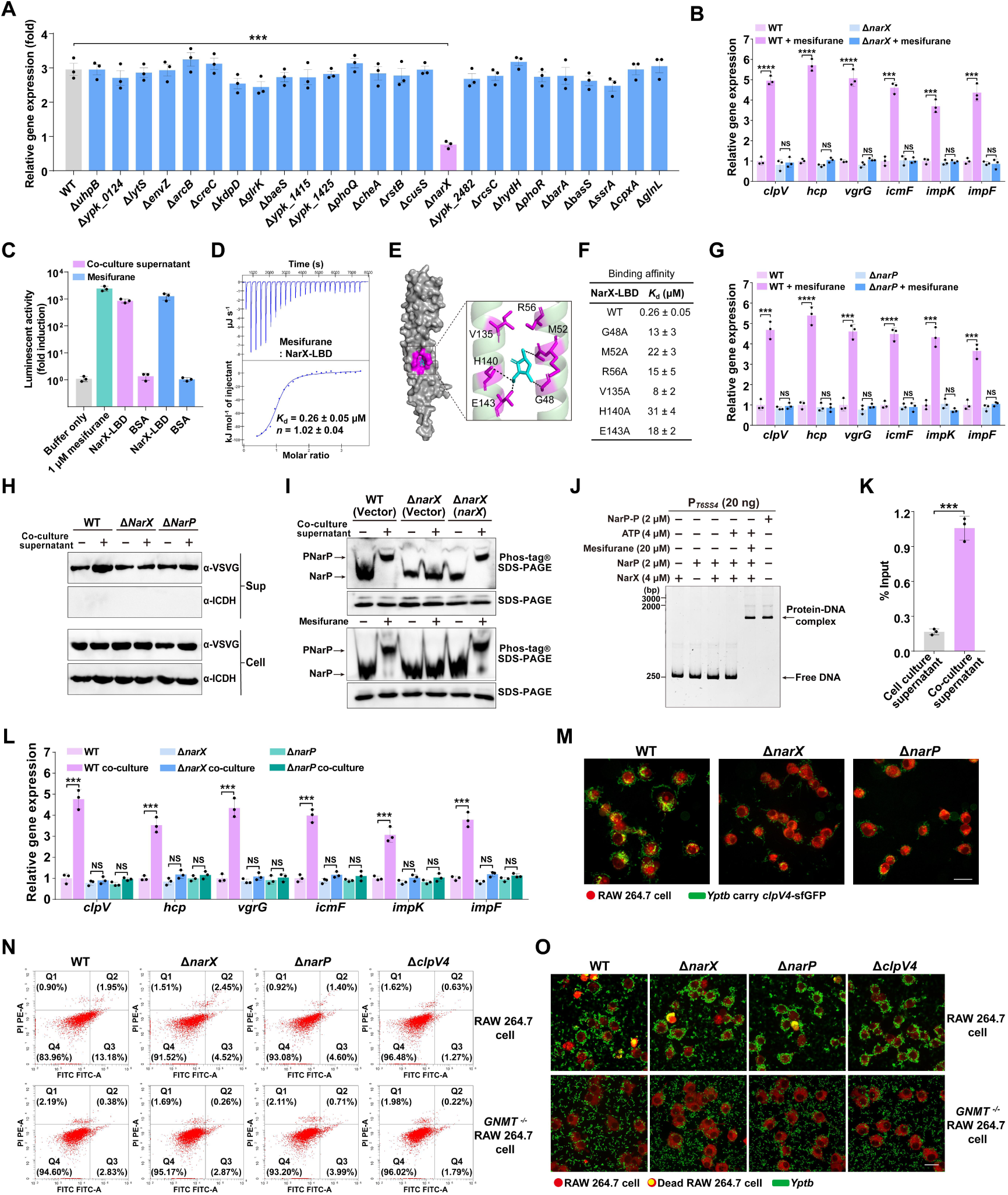
The NarX–NarP two-component system senses mesifurane to activate T6SS-4 and enhance macrophage cytotoxicity in *Yptb*. **(A)** NarX is required for mesifurane-induced T6SS transcription. *Yptb* strains harboring a chromosomal *P^T6SS4::^lacZ* reporter were grown to the exponential phase in M9 medium, then treated with co-culture supernatant for 2 h. Promoter activity was compared between strains treated with co-culture supernatant and the same strains treated with cell culture supernatant (control). Data are fold change relative to the respective control. **(B)** qRT-PCR analysis of T6SS core gene expression in WT and Δ*narX Yptb* left untreated or stimulated with 10 μM mesifurane for 2 h. Expression was normalized to 16S rRNA and is presented as fold change relative to the untreated WT *Yptb* control. **(C)** NarX directly binds the mesifurane. Purified NarX-LBD was incubated with co- culture supernatant or mesifurane; the captured ligand was heat-eluted and quantified using the *V. harveyi* MM32 bioluminescence reporter assay. 1 μM mesifurane served as a positive control, and BSA served as a negative control. Data are fold induction relative to the buffer control. **(D)** ITC analysis of the binding affinity between NarX-LBD and mesifurane. The displayed thermogram is one representative of three independent experiments; *K*^d^ and binding stoichiometry (*n*) are presented as mean ± s.d. from three independent experiments. **(E)** Predicted binding mode of mesifurane within the NarX-LBD. The optimal docking conformation of mesifurane (cyan sticks) was determined using AutoDock4.42 and Chimera. Key interacting residues are shown as purple sticks, with dashed lines indicating potential hydrogen bonds. **(F)** Binding of mesifurane to NarX-LBD and its mutants. The binding affinity was measured by ITC. The *K*^d^ values are presented as mean ± s.d. of three independent experiments. **(G)** qRT-PCR analysis of T6SS core gene expression in WT and Δ*narP Yptb* left untreated or stimulated with 10 μM mesifurane for 2 h. Expression was normalized to 16S rRNA and is presented as fold change relative to the untreated WT *Yptb* control. **(H)** The mesifurane enhances T6SS secretion in a NarX- and NarP-dependent manner. Western blot analysis of Hcp-VSVG in cell pellets (Cell) and concentrated culture supernatants (Sup) from indicated *Yptb* strains expressing Hcp-VSVG. Isocitrate dehydrogenase (ICDH) served as a loading control. The blot is representative of three independent experiments. **(I)** Mesifurane promotes NarP phosphorylation via NarX. *Yptb* strains expressing pKT100-*narP*-His^6^ were treated with co-culture supernatant or mesifurane for 2 h. Phosphorylation of NarP was analyzed by Phos-tag and conventional SDS-PAGE, followed by immunoblotting with an anti-His antibody. Blots are representative of three experiments. **(J)** Mesifurane promotes NarP binding to the T6SS-4 promoter DNA. EMSA showing the binding of NarP to a T6SS-4 promoter probe in the presence or absence of mesifurane and/or NarX. A representative gel from three independent experiments is shown. **(K)** ChIP-qPCR quantifying binding of FLAG-NarP at the promoter of the T6SS-4 gene in *Yptb* WT stimulated by co-culture supernatant for 2 h. *Yptb* treated with cell culture supernatant served as a control. The ChIP-qPCR signals were normalized to their respective DNA inputs. **(L)** T6SS gene expression during macrophage infection requires NarX and NarP. qRT-PCR analysis of T6SS gene expression in the indicated strains following infection of RAW 264.7 macrophages. Bacteria were added to macrophages at an MOI of 100, and after 2 h of co-culture, bacterial RNA was extracted for analysis. Expression was normalized to 16S rRNA and is presented as fold change relative to WT *Yptb* cultured without macrophages. **(M)** T6SS activity in macrophage co-culture requires NarX and NarP. Confocal micrographs of WT *Yptb*, Δ*narX*, and Δ*narP* strains expressing a chromosomal ClpV4-sfGFP fusion (green) following co-culture with RAW 264.7 macrophages (CellTracker Deep Red, red) at MOI 100 for 2 h. ClpV4 is the T6SS sheath- dismantling ATPase, and its focal accumulation indicates sites of active T6SS assembly. Scale bar, 20 μm. Images are representative of three independent experiments. **(N)** Flow cytometric analysis of macrophage apoptosis. WT and *GNMT*^-/-^ RAW 264.7 cells were infected with the indicated *Yptb* strains (MOI 100) for 2 h and stained with Annexin V-FITC/PI. A representative result from three independent experiments is shown. **(O)** Confocal microscopy assessment of macrophage death. Confocal micrographs of WT and *GNMT*^-/-^ RAW 264.7 cells (CellTracker™ Deep Red, red) were infected with GFP-tagged *Yptb* strains (green) at MOI 100 for 2 h. Dead macrophages were stained with PI (yellow). Scale bar, 20 μm. Images are representative of three independent experiments with similar results. Data are represented as mean ± SD of three biological replicates, each with three technical replicates. Statistical significance was determined with the two-tailed unpaired Student’s *t*-test. \*\*\**P* < 0.001; \*\*\*\**P* < 0.0001; NS, not significant.

Strikingly, we found that NarX is not just a downstream signal relay but is itself a direct receptor for host cues. Ligand capture and ITC assays confirmed that the purified NarX-LBD binds to mesifurane with high affinity (Figures 4C, 4D and S4C). Molecular docking simulations revealed the binding pocket, involving residues G48, M52, R56, V135, H140, and E143 (Figure 4E). Alanine substitution at these sites progressively weakened binding, with a triple mutation (M52A/H140A/E143A) completely abolishing it (Figures 4F, S4D and S4E).

We next asked how mesifurane binding to NarX is relayed to activate T6SS-4 transcription. In *E. coli*, NarX pairs with the response regulators NarL and NarP^40^. Genomic analysis revealed *narP* (*ypk_1384*) but not *narL* in *Yptb*, suggesting a dedicated NarX-NarP system. Accordingly, the induction of T6SS-4 genes by mesifurane was abrogated in Δ*narP* (Figures 4G and S4F). Functionally, secretion of the T6SS effector Hcp, a hallmark of active T6SS assembly^41^, was stimulated in the WT but not in the Δ*narX* or Δ*narP* strains (Figure 4H). Mesifurane stimulated NarX- dependent phosphorylation of NarP in vivo and in vitro (Figures 4I and S4G). Phosphorylated NarP (NarP∼P) bound specifically to the T6SS-4 promoter according to an electrophoretic mobility shift assay (EMSA) (Figure S4H), and a fully reconstituted in vitro phosphorelay confirmed that mesifurane initiates signaling, culminating in promoter binding (Figures 4J and S4I). Chromatin immunoprecipitation followed by qPCR (ChIP‒qPCR) confirmed increased NarP occupancy at the T6SS-4 promoter upon mesifurane stimulation in vivo (Figure 4K). These data establish a direct signaling pathway in which mesifurane is sensed by NarX, leading to NarP phosphorylation and subsequent transcriptional activation of T6SS-4.

We next examined the functional importance of this NarX–NarP axis during infection. T6SS-4 gene induction and assembly upon macrophage contact, as measured by qRT-PCR and the ClpV4-sfGFP reporter, were absent in the Δ*narX* and Δ*narP* mutants (Figures 4L and 4M). Consequently, these mutants, like the Δ*clpV4* mutant, were severely attenuated in their ability to induce macrophage apoptosis and death (Figures 4N and 4O). This defect was mirrored when *GNMT^-/-^* macrophages were infected with WT bacteria (Figures 4N and 4O), demonstrating that host mesifurane production is required for NarX-NarP-dependent T6SS-4 activation and macrophage killing.

Key mesifurane-binding residues in NarX are highly conserved across many pathogens (Figures S5A and S5B). Purified NarX homologs from *Yersinia pestis*, *Pseudomonas aeruginosa*, *Vibrio parahaemolyticus*, *Salmonella* Typhimurium, *Escherichia coli*, and even environmental bacteria all bound mesifurane with high affinity (Figures S5C-S5E). Heterologous expression of these foreign *narX* genes in the *Yptb* Δ*narX* mutant restored T6SS-4 induction (Figures S5F-S5H). Most critically, we fully reconstituted the pathway in the plague agent *Y. pestis*, showing that mesifurane stimulates *Y. pestis* NarX autokinase activity, phosphotransfer to NarP, and NarP∼P binding to its cognate T6SS promoter (Figures S5I-S5L). These findings suggest that co-opting the NarX–NarP system to sense host metabolites and directly activate virulence is a broadly conserved strategy among bacterial pathogens.

### Mesifurane relieves CusR-mediated repression of bacterial AI-2 production

Our transcriptomic data revealed another layer of regulation: mesifurane strongly induced the expression of *luxS*, the bacterial gene encoding AI-2 synthase. This suggested that host cues might amplify the pathogen’s own signaling. We confirmed this induction by qRT‒PCR (Figure S6A). We next sought to identify the regulator responsible for this induction. This led us to CusR (*YPK_2192*), an OmpR-family transcriptional repressor and a functional homolog of the *Campylobacter jejuni* LuxS repressor CosR^42^. Deletion of *cusR* derepressed *luxS* expression, and mesifurane failed to further induce *luxS* in this background (Figure 5A). A *luxS* promoter-reporter assay yielded consistent results (Figure S6B), indicating that mesifurane acts by relieving CusR-mediated repression.

**Figure 5.**
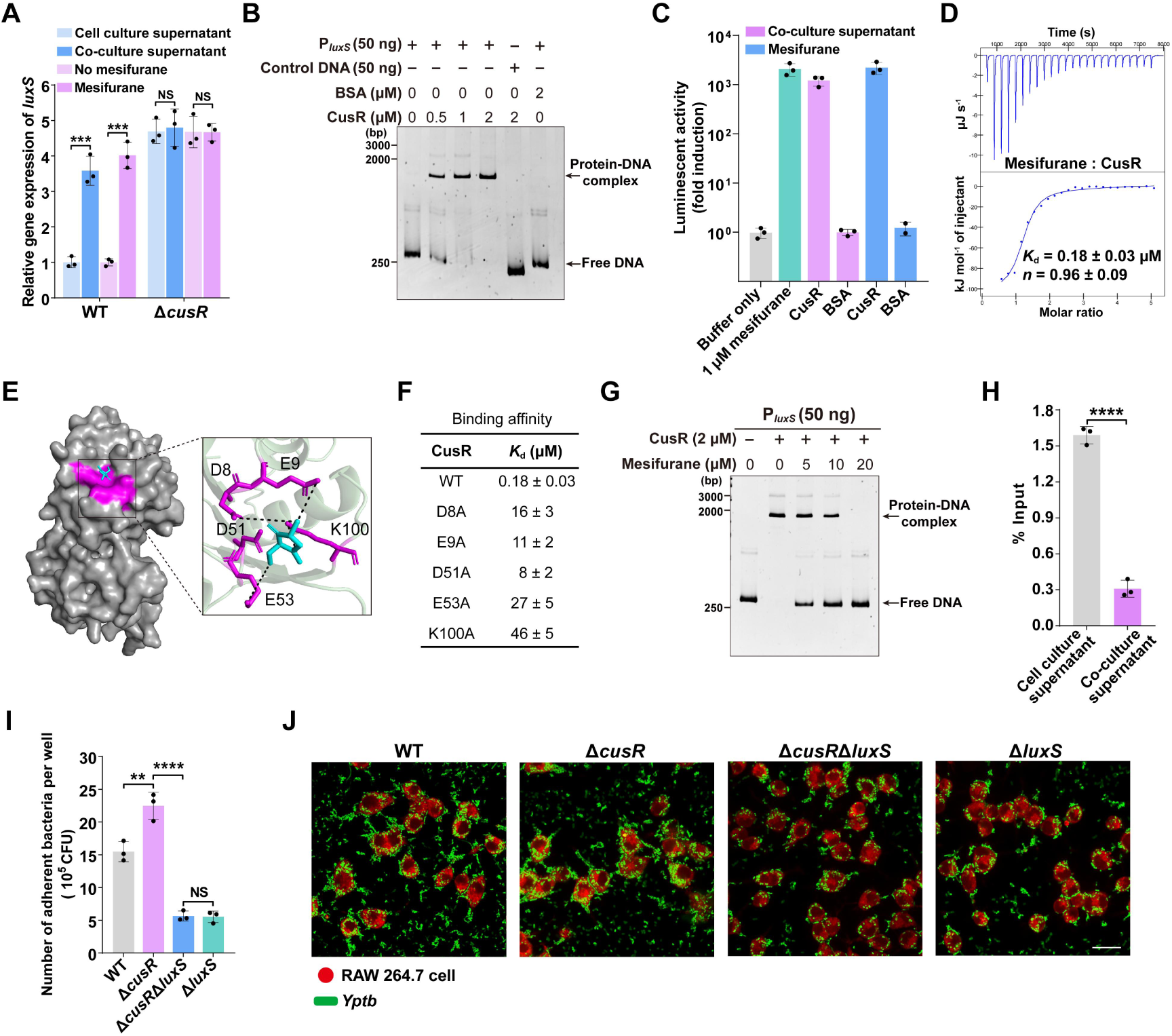
Mesifurane binds CusR to derepress *luxS* and amplify AI-2 synthesis in *Yptb*. **(A)** qRT-PCR of the *luxS* gene in WT and Δ*cusR* strains after 2 h of stimulation. Cells were treated with cell culture supernatant (unstimulated control), co-culture supernatant (containing mesifurane), or 10 μM mesifurane. Expression was normalized to 16S rRNA and is presented as fold change relative to the WT unstimulated control. **(B)** CusR binds the *luxS* promoter in vitro. EMSA showing CusR binding to the *luxS* promoter region. BSA and DNA fragments from the *luxS* coding region served as negative controls. A representative gel from three experiments is shown. **(C)** CusR interacts with the mesifurane. Purified CusR was incubated with co-culture supernatant or mesifurane; the captured ligand was heat-eluted and quantified via the *V. harveyi* MM32 bioluminescence assay. 1 μM mesifurane served as a positive control, and BSA served as a negative control. Data are fold induction relative to the buffer control. **(D)** ITC analysis of the binding affinity between CusR and mesifurane. The displayed thermogram is one representative of three independent experiments; *K*^d^ and binding stoichiometry (*n*) are presented as mean ± s.d. from three independent experiments. **(E)** Predicted binding mode of mesifurane within the CusR ligand-binding pocket. The optimal docking conformation of mesifurane (cyan sticks) was determined using AutoDock4.42 and Chimera. Key interacting residues are shown as purple sticks, with dashed lines indicating potential hydrogen bonds. **(F)** Binding of mesifurane to CusR and its mutants. The binding affinity was measured by ITC. *K*^d^ values are presented as mean ± s.d from three experiments. **(G)** Mesifurane reduces CusR binding to the *luxS* promoter DNA. EMSA for CusR binding to the *luxS* promoter in the presence or absence of mesifurane. Mesifurane was simultaneously added with CusR to the reaction system. A representative gel from three experiments is shown. **(H)** ChIP-qPCR quantifying binding of FLAG-CusR at the promoters of the *luxS* genes in *Yptb* WT stimulated by co-culture supernatant for 2 h. *Yptb* treated with cell culture supernatant served as a control. The ChIP-qPCR signals were normalized to their respective DNA inputs. **(I)** Quantification of *Yptb* adherence to macrophages. RAW 264.7 macrophages were infected with the indicated *Yptb* strains at an MOI of 100 for 1 h. Nonadherent bacteria were removed by washing, and adherent bacteria were released by cell lysis, serially diluted, and enumerated by CFU plating. **(J)** RAW 264.7 macrophages (CellTracker™ Deep Red, red) were infected with GFP- expressing *Yptb* strains (green) at MOI 100 for 1 h. Nonadherent bacteria were removed by washing prior to imaging. Scale bar, 20 μm. Images are representative of three independent experiments. Data are represented as mean ± SD of three biological replicates, each with three technical replicates. Statistical significance was determined with the two-tailed unpaired Student’s *t*-test. \*\**P* < 0.01; \*\*\**P* < 0.001; \*\*\*\**P* < 0.0001; NS, not significant.

We next asked whether mesifurane directly regulates CusR activity. EMSAs confirmed that CusR binds specifically to the *luxS* promoter region (Figure 5B). Crucially, ligand capture and ITC assays demonstrated that mesifurane binds directly to CusR with high affinity (Figures 5C, 5D and S6C). Docking positioned mesifurane in a CusR pocket involving residues D8, E9, D51, E53, and K100; mutagenesis of these sites impaired binding (Figures 5E, 5F and S6D). Functionally, increasing concentrations of mesifurane disrupted the formation of CusR-*luxS* promoter complexes according to EMSA (Figures 5G and S6E), and ChIP‒qPCR revealed that stimulation with mesifurane reduced CusR occupancy at the *luxS* promoter in vivo (Figure 5H). Thus, the host metabolite directly antagonizes a bacterial repressor to derepress AI-2 production.

Mesifurane-mediated derepression of *luxS* creates a powerful positive feedback loop. The Δ*cusR* mutant, with constitutively high AI-2 production, exhibited enhanced adhesion to macrophages (Figures 5I and 5J). However, a Δ*cusR*Δ*luxS* double mutant, which cannot produce AI-2, showed a ∼4-fold reduction in adhesion relative to the Δ*cusR* single mutant, phenocopying the Δ*luxS* single mutant (Figures 5I and 5J). This finding demonstrates that the adhesion advantage gained from CusR derepression is entirely mediated by the resulting increase in endogenous AI-2. The host signal thus not only guides individual bacteria but also triggers them to broadcast an amplified AI-2 signal to the surrounding population, promoting collective recruitment. This regulatory module is conserved in *Y. pestis* (Figures S6F-S6I).

### An integrated “sense, arm, recruit” circuit drives pathogenesis and is a therapeutic target

Collectively, our findings define a coordinated, three-module circuit through which a single host metabolite orchestrates bacterial virulence. Mesifurane engages three dedicated sensory systems: YcrA, which drives chemotaxis and adhesion (Sense); NarX-NarP, which licenses T6SS-mediated killing (Arm); and CusR, whose antagonism amplifies endogenous AI-2 production to recruit additional bacteria (Recruit).

We next asked whether this integrated circuit operates in intestinal epithelial cells, a physiologically relevant barrier that pathogens must breach^43,44^. During infection of Caco-2 intestinal epithelial cells, the entire mesifurane-sensing network was similarly operational. Caco-2 cells produced mesifurane during coculture with bacteria (Figures 2J-2L), and adhesion was significantly impaired in the Δ*ycrA* mutant, while Δ*cusR* exhibited enhanced attachment (Figures S7A and S7B). Both the Δ*luxS* and Δ*cusR*Δ*luxS* double mutants showed severe adhesion defects (Figures S7A and S7B), mirroring the adhesion phenotype observed in macrophages. During Caco-2 infection, the expression of T6SS-4 core genes was strongly induced in WT *Yptb*, but this response was abolished in the Δ*narX* and Δ*narP* mutants (Figure S7C). Accordingly, epithelial apoptosis was significantly attenuated upon infection with Δ*narX*, Δ*narP*, or the T6SS structural mutant Δ*clpV4* (Figure S7D). These data confirm that the entire mesifurane-sensing network, from chemotaxis and adhesion to T6SS activation, is functional within intestinal epithelial cells, indicating that this circuit is not restricted to professional phagocytes.

To assess the contribution of this network to virulence in vivo, we first examined lethality in a mouse infection model. Kaplan‒Meier survival analysis revealed that intragastric infection of mice with WT *Yptb* caused more than 80% mortality within three weeks, whereas infection with Δ*ycrA,* Δ*narX*, or Δ*ycrA*Δ*narX* resulted in significantly reduced mortality (Figure 6A). A parallel systemic mouse infection model (intraperitoneal challenge) confirmed that these mutants also markedly reduced mortality (Figure 6B).

**Figure 6.**
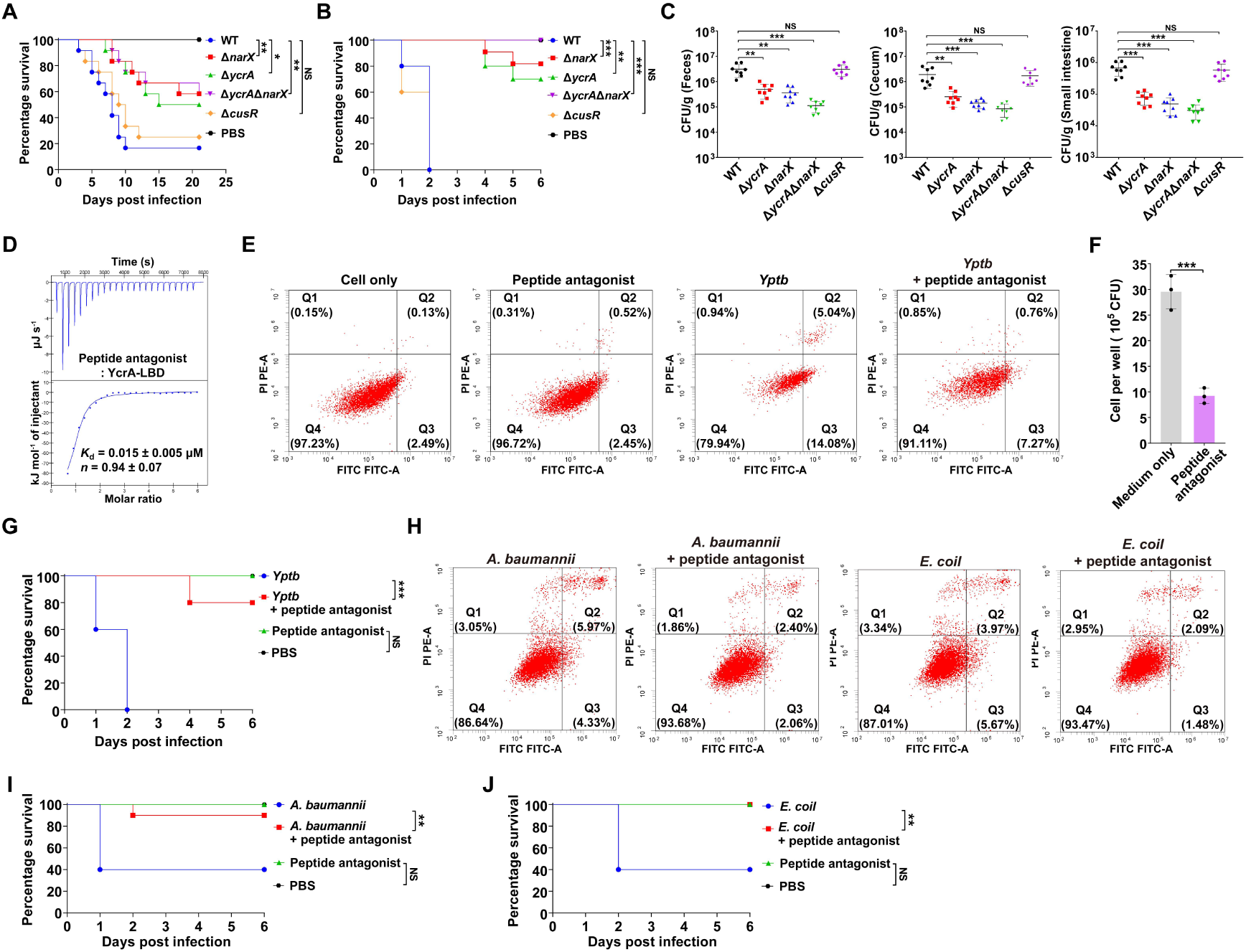
A Peptide Antagonist Attenuates Virulence of Multidrug-Resistant Clinical Isolates. (A) Survival curves of BALB/c mice following oral infection with indicated *Yptb* strains (1 × 10^9^ CFU) (n = 12 mice per group). (B) Survival curves of BALB/c mice following intraperitoneal infection with indicated *Yptb* strains (1 × 10^7^ CFU) (n = 10 mice per group). (C) The mesifurane signaling enhances *Yptb* colonization in vivo. Bacterial burdens in feces, cecum, and intestinal tissues of BALB/c mice at 2 days post-intragastric infection with relevant *Yptb* strains (1 × 10^9^ CFU) (n = 8 mice per group). (D) The peptide antagonist binds YcrA with high affinity. ITC analysis of the interaction between YcrA and the peptide antagonist. The displayed thermogram is one representative of three independent experiments; *K*^d^ and binding stoichiometry (*n*) are presented as mean ± s.d. from three independent experiments. (E) Peptide antagonist attenuates *Yptb*-induced macrophage apoptosis. RAW 264.7 cells were left untreated, treated with peptide antagonist (5 μg/ml) alone, infected with WT *Yptb* (MOI 100) alone, or simultaneously exposed to peptide antagonist (5 μg/ml) and WT *Yptb*. Apoptosis was assessed 2 h later by Annexin V-FITC/PI staining. Representative flow cytometry plots from three independent experiments are shown. (F) The peptide antagonist inhibits *Yptb* chemotaxis toward macrophages. Quantification of WT *Yptb* migration toward RAW 264.7 macrophages in a transwell assay, in the presence or absence of 5 µg/ml peptide antagonist. Final CFU counts were corrected by subtracting the number of bacteria that migrated into cell-free control wells. (G) Peptide antagonist treatment in a murine model of *Yptb* infection. Mice were infected intraperitoneally with WT *Yptb* (1 × 10^7^ CFU) and simultaneously treated with peptide antagonist (10 mg/kg) or PBS. Uninfected controls received peptide antagonist or PBS alone. Treatment was repeated every 24 h for 6 days. Survival was monitored for 6 days. (n = 10 mice per group). (H) The peptide antagonist reduces macrophage apoptosis induced by bacterial infection. RAW 264.7 cells infected with the indicated clinical isolates (MOI 100) alone, or simultaneously exposed to 5 µg/ml peptide antagonist and bacterial strains. Apoptosis was assessed 2 h later by Annexin V-FITC/PI staining. A representative result from three independent experiments is shown. **(I and J)** Mice were infected intraperitoneally with *A. baumannii* Ab-C17 (5 × 10^5^ CFU) (I) or *E. coli* EC155 (1 × 10^7^ CFU) (J) and simultaneously treated with peptide antagonist (10 mg/kg) or PBS. Uninfected controls received peptide antagonist or PBS alone. Treatment was repeated every 24 h for 6 days. Survival was monitored for 6 days. (n = 10 mice per group). For mouse survival assays (A, B, G, I, J), two independent experiments were performed; one representative experiment is shown, with the indicated number of mice per group. For tissue bacterial burden (C), data are presented as mean ± SD from eight biological replicates (individual mice), each assessed in three technical replicates. For (F), data are presented as mean ± SD of three biological replicates, each with three technical replicates. Statistical significance was determined with the two-tailed unpaired Student’s *t*-test (C, F) or Log-rank (Mantel–Cox) test (A, B, G, I, J). \**P* < 0.1; \*\**P* < 0.01; \*\*\**P* < 0.001; \*\*\*\**P* < 0.0001; NS, not significant.

To test whether host-derived mesifurane alone, in the absence of endogenous AI-2, is sufficient to cause lethal infection in mice, we compared survival following intraperitoneal challenge with the WT, Δ*luxS*, Δ*luxS*Δ*ycrA*, and Δ*luxS*Δ*narX* strains. The Δ*luxS* mutant was as lethal as the WT, with both groups succumbing within two days, indicating that endogenous AI-2, while contributing to adhesion in vitro, is not required for full virulence in vivo. In contrast, disruption of either mesifurane sensing (Δ*luxS*Δ*ycrA*) or T6SS licensing (Δ*luxS*Δ*narX*) in the AI-2-deficient background significantly reduced mortality (Figure S7E).

To determine whether these pathways contribute to intestinal colonization, we quantified bacterial burdens in the feces, cecum, and intestine 48 h after infection. Bacterial loads were significantly reduced in mice infected with the Δ*ycrA,* Δ*narX*, or Δ*ycrA*Δ*narX* double mutant compared with those in WT controls (Figure 6C), demonstrating that both YcrA-mediated chemotaxis/adhesion and NarX-mediated T6SS activation are critical for efficient intestinal colonization. The Δ*cusR* mutant colonized as effectively as the WT did (Figure 6C), indicating that CusR-dependent AI-2 amplification is unnecessary for intestinal colonization. Combined with the finding that endogenous AI-2 is dispensable for lethal infection (Figure S7E), these data establish the CusR module as an accessory amplifier rather than a core virulence determinant. Thus, among the three sensory modules, YcrA-driven chemotaxis and NarX-mediated T6SS activation constitute the core virulence program.

The indispensable role of YcrA in virulence, combined with its well-defined ligand-binding pocket, suggested a therapeutic opportunity: blocking this sensor should disarm the entire circuit. To test this, we designed a panel of peptide antagonists by computationally screening the mesifurane-binding pocket, focusing on the critical residues identified above (R120, W122, Y138, D140, and D167), and refined candidates through docking-based screening and AlphaFold3 validation (Table S4). The lead candidate (sequence: GKKYYTIGKYDYEK) bound the YcrA- LBD with exceptional affinity (*K*^d^ = 0.015 ± 0.005 μM), occupying the predicted binding pocket in molecular docking models (Figures 6D and S7F). This peptide was nontoxic to mammalian cells at effective concentrations (Figure 6E), blocked *Yptb* chemotaxis toward macrophages (Figure 6F) and significantly reduced macrophage apoptosis induced by infection in vitro (Figure 6E). Most importantly, this YcrA antagonist had dramatic efficacy in vivo. In a lethal systemic infection model, the administration of the peptide at the time of infection and every 24 h thereafter resulted in a significant survival advantage, whereas all the untreated mice died within two days (Figure 6G). These findings demonstrate that pharmacologic inhibition of YcrA with a peptide antagonist is sufficient to disrupt the mesifurane- sensing circuit and attenuate *Yptb* virulence in vivo.

Given the high sequence conservation of the YcrA-LBD across multiple Gram- negative pathogens, including carbapenem-resistant clinical isolates of *E. coli* (EC155)^45^, *A. baumannii* (Ab-C17, ST457)^46^, and *P. aeruginosa* (ATCC BAA-2108)^47^ (Figure S7G), we next tested whether this peptide antagonist could also attenuate infection by these clinically relevant resistant strains. The lead YcrA antagonist potently attenuated the cytotoxicity of *E. coli* (EC155) and *A. baumannii* (Ab-C17 ST457) toward macrophages (Figure 6H) and significantly improved mouse survival rates in systemic infection models. (Figures 6I and 6J).

These results establish YcrA as a druggable target and demonstrate that blocking this sensor with a peptide antagonist is sufficient to disrupt the entire virulence circuit, not only in the model pathogen but also in multidrug-resistant clinical isolates. Given the therapeutic promise of this approach, we next asked whether clinically approved drugs could similarly target YcrA. Suramin sodium, a small molecule with an established safety profile, emerged as a potent inhibitor of YcrA- mediated virulence in vivo.

### Suramin sodium targets YcrA to disarm multidrug-resistant pathogens in vivo

Proof-of-concept peptide studies have established YcrA as a druggable target. To identify small-molecule inhibitors with greater translational potential, we performed structure-based virtual screening of a library of ∼26,000 clinically approved and investigational compounds against the mesifurane binding pocket of YcrA, from which we selected the highest-affinity candidates (docking score ≤ −9.0 kcal mol⁻^1^) (Table S5). Among them, suramin sodium, a century-old antiparasitic drug with a well-characterized safety profile, emerged as the lead^26,27,48^, offering a unique drug-repurposing opportunity.

We first confirmed that suramin sodium directly binds YcrA. ITC revealed that suramin sodium bound the recombinant YcrA-LBDs from all three pathogens, *E. coli* (EC155), *A. baumannii* (Ab-C17, ST457), and *P. aeruginosa* (ATCC BAA-2108), with dissociation constants in the extremely low range (Figure 7A). Molecular docking using the crystal structure of *P. aeruginosa* YcrA revealed that suramin sodium occupies the entire mesifurane binding pocket, sterically precluding ligand access (Figure S8A).

**Figure 7.**
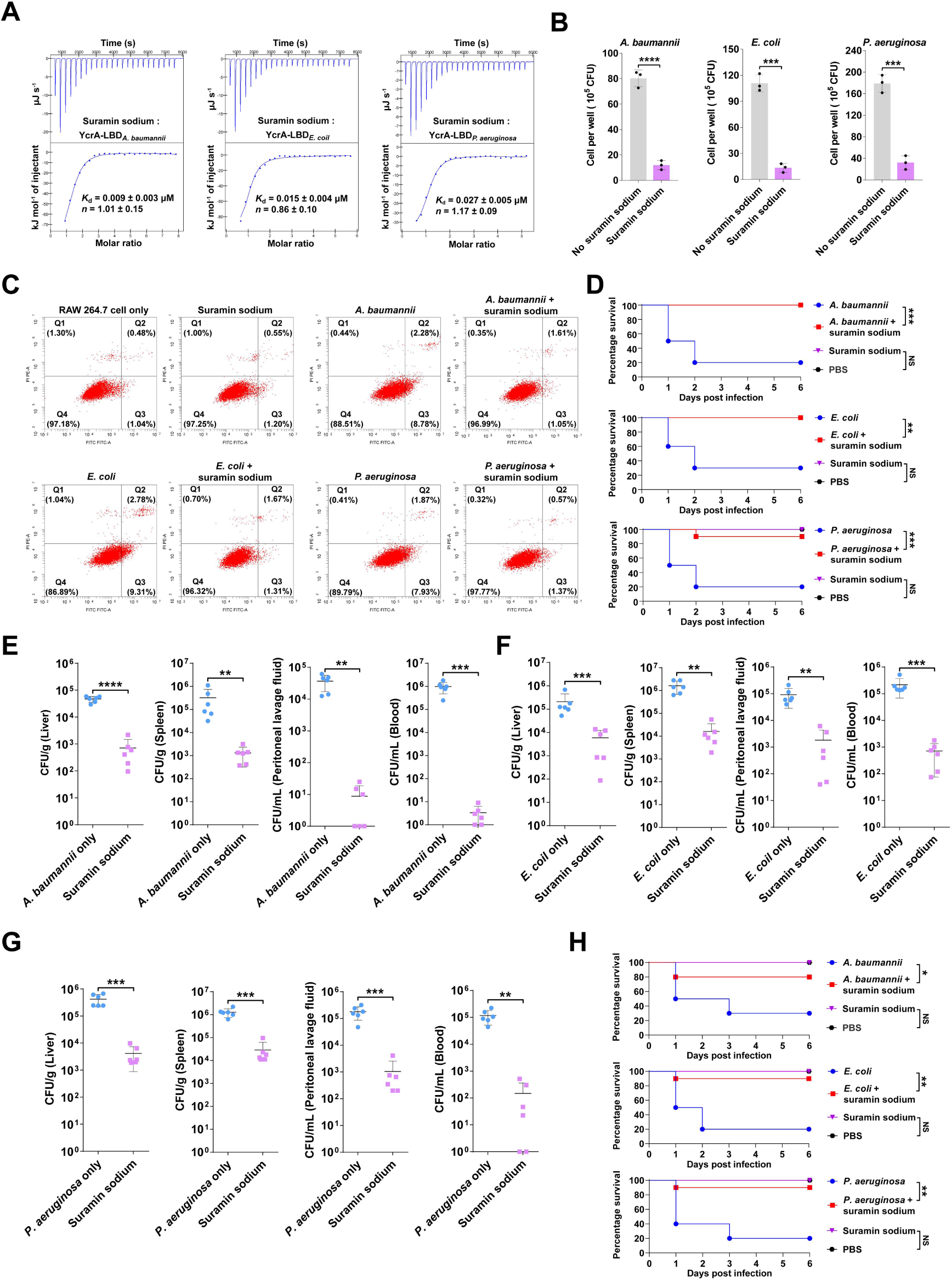
Suramin sodium blocks YcrA-dependent chemotaxis and protects mice from lethal infection. **(A)** ITC analysis of suramin sodium binding to recombinant YcrA-LBDs from *A. baumannii* Ab-C17, *E. coli* EC155, and *P. aeruginosa* ATCC BAA-2108. The displayed thermogram is one representative of three independent experiments; *K*^d^ and binding stoichiometry (*n*) are presented as mean ± s.d. from three independent experiments. **(B)** Suramin sodium inhibits the indicated clinical isolates’ chemotaxis toward macrophages. Quantification of the indicated clinical isolates’ migration toward RAW 264.7 macrophages in a transwell assay, in the presence or absence of 5 µg/ml suramin sodium. Final CFU counts were corrected by subtracting the number of bacteria that migrated into cell-free control wells. **(C)** Suramin sodium reduces macrophage apoptosis induced by bacterial infection. RAW 264.7 cells were left untreated, treated with suramin sodium (5 μg/ml) alone, infected with the indicated bacterial strains (MOI 100) alone, or simultaneously exposed to suramin sodium and the indicated bacterial strains. Apoptosis was assessed 2 h later by Annexin V-FITC/PI staining. Representative flow cytometry plots from three independent experiments are shown. **(D)** Delayed suramin sodium treatment rescues mice from lethal infection with the indicated clinical isolates. Survival of mice infected intraperitoneally with *A. baumannii* Ab-C17 (1× 10^6^ CFU), *E. coli* EC155 (2 × 10^7^ CFU), or *P. aeruginosa* ATCC BAA-2108 (5× 10^6^ CFU). Infected mice received suramin sodium (10 mg/kg) or PBS subcutaneously beginning 2 h post-infection, every 12 h for the first 3 days and every 24 h thereafter. Uninfected controls were given suramin sodium or PBS on the same schedule. Survival was monitored for 6 days. (n = 10 mice per group). **(E-G)** Suramin sodium reduces bacterial dissemination across infection models of carbapenem-resistant pathogens. Bacterial loads in blood, peritoneal lavage fluid, spleen, and liver from mice infected with (E) *A. baumannii* Ab-C17 (1× 10^6^ CFU), (F) *E. coli* EC155 (2 × 10^7^ CFU), or (G) *P. aeruginosa* ATCC BAA-2108 (5× 10^6^ CFU). Infected mice received suramin sodium (10 mg/kg) or PBS subcutaneously beginning 2 h post-infection, and tissues were collected 14 h post-infection. (n = 6 mice per group). Data are presented as CFU per ml (blood, lavage) or CFU per g (tissue). **(H)** Suramin sodium treatment initiated at 8 h post-infection also confers protection against lethal infection. Survival of mice infected and treated as described in (D), except that suramin sodium (10 mg/kg) or PBS was administered subcutaneously beginning 8 h post-infection instead of 2 h. Survival was monitored for 6 days. (n = 10 mice per group). For mouse survival assays (D, H), two independent experiments were performed; one representative experiment is shown, with the indicated number of mice per group. For tissue bacterial burden (E–G), data are presented as mean ± SD from six biological replicates (individual mice), each assessed in three technical replicates. For (B), data are presented as mean ± SD of three biological replicates, each with three technical replicates. Statistical significance was determined with the two-tailed unpaired Student’s *t*-test (B, E–G) or Log-rank (Mantel–Cox) test (D, H). \**P* < 0.1; \*\**P* < 0.01; \*\*\**P* < 0.001; \*\*\*\**P* < 0.0001; NS, not significant.

We next tested whether suramin sodium could functionally block mesifurane- driven chemotaxis and cytotoxicity. In Transwell assays, suramin sodium significantly reduced the chemotaxis of all three clinical isolates toward macrophages (Figure 7B). Similarly, the drug markedly decreased macrophage apoptosis induced by each pathogen, with no detectable cytotoxicity toward mammalian cells at the concentrations tested (Figure 7C). Suramin sodium did not inhibit bacterial growth at concentrations up to 200 μM, far exceeding those required for cellular efficacy (Figure S8B). Thus, suramin sodium disarms pathogen virulence without directly impairing bacterial viability.

We then evaluated the therapeutic efficacy of suramin sodium in vivo. Mice were infected intraperitoneally with each of the three multidrug-resistant bacterial isolates and treated with suramin sodium beginning 2 h post infection. This regimen conferred striking protection: survival rates reached 100% for *E. coli* (EC155) and *A. baumannii* (Ab-C17, ST457) and 90% for *P. aeruginosa* (ATCC BAA-2108), compared with 20– 30% in PBS-treated controls (Figure 7D). Consistent with the improved survival, bacterial burdens in the blood, peritoneal lavage fluid, spleen, and liver were significantly reduced in suramin sodium-treated mice compared to controls across all three infection models at 12 h post treatment (Figures 7E–7G).

Systemic infection often triggers a dysregulated inflammatory response that contributes to lethality^49,50^. Suramin sodium treatment significantly reduced the levels of IL-1β, IL-6, IL-10, TNF-α, and IFN-γ in the serum, peritoneal lavage fluid, spleen, and liver in all three infection models (Figures S8C–S8E), with corresponding suppression of cytokine transcript levels (Figures S9A–S9C).

Serial passage of all three pathogens for approximately 200 generations in the presence of suramin sodium did not lead to the emergence of resistant mutants; passaged bacteria remained fully virulent in untreated mice and retained complete sensitivity to suramin sodium treatment in vivo (Figure S9D). Thus, suramin sodium did not select for resistant mutants under the conditions tested, highlighting the reduced selective pressure associated with targeting host-directed chemotaxis rather than bacterial viability.

Finally, we tested whether suramin sodium could rescue mice when administered at a clinically realistic time point after infection onset. Even when the first dose was delayed until 8 h post infection, a time at which severe infection is already established, approximately 90% of mice survived across all three pathogens (Figure 7H), demonstrating a broad therapeutic window.

Collectively, our findings delineate a complete pathogenic circuit driven by a single host metabolite and establish YcrA as a therapeutically actionable target. The repurposing of suramin sodium provides a direct path to clinical translation and validates a new paradigm for combating antimicrobial resistance by disarming, rather than killing, bacterial pathogens.

## Discussion

The host-pathogen interface is shaped by a continuous exchange of chemical signals. Bacteria sense a range of host-derived cues^9,51^, including nutrients^52^, hormones^10^, and immune effectors^53^, to gauge their environment and adjust virulence accordingly. However, whether host-derived cues can function as master regulators that simultaneously coordinate disparate virulence functions, rather than merely triggering individual responses, has remained unexplored. Here, we reveal how a single host metabolite can be co-opted by a bacterial pathogen to orchestrate a complete offensive strategy, and identify this host-sensory circuit as a therapeutically targetable vulnerability. We demonstrate that *Yptb* intercepts the host-derived signal mesifurane through three dedicated sensory systems to execute a coordinated “Sense, Arm, Recruit” program: YcrA-driven chemotaxis toward host cells, NarX- NarP-mediated T6SS killing, and CusR-mediated amplification of endogenous AI-2 to recruit additional bacteria. Pharmacological blockade of the central sensor YcrA with suramin sodium disarms this entire circuit, converting near-uniform lethality to near- complete survival in models of infection with carbapenem-resistant *E. coli*, *A. baumannii*, and *P. aeruginosa*. These findings establish pathogen sensory inhibition as an anti-infective strategy that disarms rather than kills, offering a treatment path that circumvents conventional antibiotic resistance mechanisms (Figure S10).

This study identifies the host-derived chemoattractant as mesifurane, a furanone derivative. We traced its biosynthetic origin to a GNMT-dependent methyltransferase reaction. Concurrently, another group recently identified l-xylosone, a sugar-derived α-diketone, as a chemically distinct AI-2 mimic from Caco-2 cells^25^. The two molecules differ in structure, biosynthetic origin, and borate dependence, suggesting that mammalian cells may deploy multiple AI-2 mimics as part of a broader chemical lexicon for host-microbe communication. Whether different mimics activate distinct bacterial programs or function in different infection contexts remains an open question.

Crucially, our work reveals that mesifurane is not merely a chemical signal. Its production is triggered by a secreted bacterial factor at exceedingly low doses (Figure S1K), ruling out nonspecific cellular stress as the inducer and supporting its role as a specific host alarm cue. The pathogen has evolved to surveil and interpret this host broadcast as a reliable cue to launch an attack, effectively turning a host distress signal into a pathogen invasion command^9,51,54,55^.

The three sensory systems we identify—YcrA, NarX-NarP, and CusR—illustrate how canonical bacterial proteins are repurposed for host-directed warfare. The function of the chemoreceptor YcrA expands from sensing bacterial AI-2 to directly binding a host metabolite, enabling precise chemotaxis toward immune cells. The NarX-NarP two-component system, which is ancestrally tied to nitrate respiration^40^, is co-opted as a direct T6SS activator. This provides a “just-in-time” licensing mechanism, ensuring that this energetically costly weapon is deployed only when the pathogen is in immediate proximity to its target following YcrA-mediated chemotaxis.

Concurrently, by antagonizing the repressor CusR, mesifurane amplifies endogenous AI-2 production, creating a self-reinforcing recruitment loop. Importantly, this tripartite circuit is governed by a clear functional hierarchy: host-derived mesifurane, not bacterially produced AI-2, is the dominant driver.

Mesifurane outcompetes AI-2 in directing chemotaxis and selectively activates the T6SS: compared with AI-2, mesifurane elicited stronger chemotaxis at every matched concentration (Figure S2G), and in a Δ*luxS* mutant, supplementing AI-2 at concentrations exceeding its physiological peak failed to induce T6SS, whereas mesifurane at matched concentrations triggered robust expression (Figure S3D). Importantly, the Δ*luxS* mutant, which cannot produce AI-2, retained full virulence in mice, but mesifurane is sufficient to drive lethal infection in the complete absence of bacterial AI-2 (Figure S7E). AI-2, as a classical quorum-sensing signal, requires accumulation to a threshold concentration dependent on population density^19,20,56^, a constraint that inherently delays its availability during early infection^57,58,59^. Mesifurane, by contrast, is produced by host cells in direct response to a secreted bacterial trigger at exceedingly low doses (Figure S1K), providing an immediate and reliable cue independent of bacterial numbers. Notably, *P. aeruginosa* and *A. baumannii*, which lack the AI-2 synthetic gene^21,60^, were also rendered significantly less virulent upon pharmacological blockade of their respective YcrA orthologs. These findings suggest that the pathogen evolved to prioritize this host-derived signal over its own quorum-sensing molecule as the dominant regulator of virulence, ensuring that the offensive program is deployed at the earliest possible moment when host cells are present. The requirement of both YcrA and NarX-NarP for acute lethality identifies these two sensory modules as core components of the virulence program, highlighting host signal perception as a point of therapeutic vulnerability.

Because multiple virulence modules converge on the YcrA sensory node, blocking this receptor disables the entire program, akin to disrupting a signal receiver rather than disarming each weapon individually. Using a structure-based designer peptide that occludes the YcrA ligand-binding pocket, we established the feasibility of pharmacological inhibition. Virtual screening of ∼26,000 clinically approved and investigational compounds revealed suramin sodium as a potent YcrA antagonist. In vivo, suramin sodium disrupted chemotaxis, preventing subsequent T6SS-dependent death, reducing bacterial burdens and systemic inflammation, and conferring near- complete protection. The drug remained effective when administration was delayed until 8 h post infection, a time at which severe infection was already established. This efficacy likely reflects disruption of a host-pathogen signaling dialog that remains critical for virulence even as infection progresses and bacterial burdens rise. Serial passage for more than 200 generations did not result in the emergence of resistant mutants, which is consistent with the principle that targeting a signaling pathway, rather than bacterial viability, imposes minimal selective pressure^61,62^.

These findings have particular salience for the antimicrobial resistance crisis. The three pathogens in which we validated suramin sodium—carbapenem-resistant *E. coli*^45^, *A. baumannii*^46^, and *P. aeruginosa*^47^—are all classified as WHO critical- priority pathogens^28,63,64,65^. Antivirulence strategies have long been proposed as alternatives to conventional antibiotics^2,4^, but their clinical translation has been slow^5,6,66,67^, in part because few virulence targets are both broadly conserved and amenable to pharmacological blockade^6,13,68^. YcrA meets both criteria: its ligand- binding pocket is conserved across diverse Gram-negative pathogens, yet it is not required for bacterial viability, making it an ideal target for inhibitors that disarm rather than kill. Together, these findings identify YcrA as a broadly conserved antivirulence target across clinically relevant Gram-negative pathogens.

Interfering with the conserved sensory node YcrA therefore offers a single strategy that bypasses the heterogeneous resistance profiles that complicate current therapy. The reliance of *Yptb* and related pathogens on a specific host cue creates an inherent vulnerability: mutations that disrupt drug binding to YcrA would likely also compromise recognition of the native host signal, undermining the attack program of the pathogen. This evolutionary constraint, together with the relatively open architecture of chemoreceptor binding domains, makes the repurposing of approved compounds a pragmatic strategy.

Suramin sodium, a century-old antiparasitic drug with a well-characterized clinical safety profile^26,27,48^, exploits this vulnerability by blocking the host-pathogen signaling interface. Because the drug does not kill the cells, existing resistance mechanisms, carbapenemases, efflux pumps, and permeability mutations, are irrelevant. Suramin sodium has been used clinically for decades, although its known toxicity requires careful management. In our antivirulence application, suramin targets bacterial signal perception at concentrations substantially lower than those used in conventional antiparasitic therapy^27,48,69^, potentially mitigating the dose- dependent toxicity associated with long-term, high-dose regimens. This favorable therapeutic window, combined with the drug’s established safety profile, provides a strong foundation for repurposing suramin sodium as an antivirulence agent. This study provides a foundation for future pharmacokinetic and safety studies of YcrA- targeted antivirulence therapies, which could accelerate their clinical evaluation.

Several limitations of this study should be acknowledged. First, although serial passaging did not generate resistant mutants, these experiments were conducted under defined in vitro conditions. Whether sustained drug pressure in more complex in vivo environments, where spatial heterogeneity, biofilm formation, and host immune pressure coexist, could select for YcrA escape variants remains to be tested. Second, our murine infection models, while establishing proof-of-concept for suramin sodium efficacy, do not fully recapitulate human infection, particularly with respect to adaptive immunity and polymicrobial interactions. Third, the contribution of the CusR module to virulence was not evident in acute infection models, suggesting that its role may be more relevant in chronic or biofilm-associated settings, which were not examined here. Beyond these specific limitations, the physiological function of mesifurane in uninfected mammalian cells remains unknown, and the efficacy of suramin sodium across broader panels of clinical isolates warrants further evaluation. Future studies employing longitudinal infection models, humanized immune systems, and expanded pathogen panels will be important to address these gaps. We note, however, that the core finding, that YcrA-mediated host signal perception is essential for virulence, was consistently observed across multiple infection routes and pathogen species, supporting the robustness of the central conclusion.

More broadly, this work establishes a framework for developing anti-infectives that target the sensory interface between pathogen and host. By disrupting the chemical conversations that coordinate virulence, such strategies offer a path toward therapies that remain effective even as bacteria evolve resistance to conventional antibiotics—a prospect of urgent importance in the post-antibiotic era.

## MATERIALS AND METHODS

### Bacterial strains, plasmid constructions and growth conditions

Bacterial strains and plasmids used in this study are listed in Table S6. All primers used in this study were designed using Primer premier 5.0 (Premier Biosoft) and their sequences are listed in Table S7. *Yptb* strains and their derivatives were cultured in Yersinia-Luria-Bertani (YLB) broth (1% tryptone, 0.5% yeast extract, 0.5% NaCl) or M9 minimal medium (Na_2_HPO^4^, 6 g/L; KH_2_PO_4_, 3 g/L; NaCl, 0.5 g/L; NH_4_Cl, 1 g/L; MgSO_4_, 1 mM; CaCl^2^, 0.1 mM; glucose 0.4%, pH 7.0) with appropriate antibiotics at 26°C. *E. coli*, *S*. Typhimurium, *P. aeruginosa*, *A. baumannii,* and *K. pneumoniae* strains and their derivatives were grown in LB medium with appropriate antibiotics at 37°C. *V. harveyi MM32* was grown at 30°C in AB medium^70^. In-frame deletion and point mutants of *Yptb* were generated as described^71^. For genetic complementation, target genes were cloned into the plasmid pKT100 or pME6032. To express and purify soluble His^6^-tagged recombinant proteins, genes were cloned into pET-28a, and then transformed into *E. coli* BL21(DE3) host strain. For protein secretion experiments, the pKT100 derivative containing C-terminal VSVG-tagged Hcp was constructed. The DNA fragments encoding four YcrA homologs (*Yersinia pestis*, *Shinella sp. HZN7*, *Denitrificimonas caeni,* and *Stutzerimonas degradans*), three NarX homologs (*Yersinia pestis*, *Gilliamella apicola,* and *Cupriavidus taiwanensis*), and NarP, CusR homologs from *Yersinia pestis* were synthesized by Genewiz (Suzhou, China). Bacterial growth was monitored by measuring the optical density at 600 nm (OD_600_). When necessary, antibiotics were added at the following concentrations: streptomycin, nalidixic acid, 20 μg/mL; kanamycin, 50 μg/mL; ampicillin, 100 μg/mL; chloramphenicol, 20 μg/mL; tetracycline, 10 μg/mL.

### Mammalian cell culture

RAW 264.7, HEK293T, and HeLa cells were cultured in DMEM medium supplemented with 10% FBS and 1% PenStrep. THP-1 cells were cultured in RPMI 1640 medium supplemented with 10% FBS, 1% PenStrep, sodium pyruvate (10 μM), L-Glutamine (2 mM), nonessential amino acids (0.1 mM), 2-mercaptoethanol (50 μM), and HEPES (25 mM). Caco-2 cells were maintained in MEM medium supplemented with 20% FBS and 1% PenStrep. All cell lines were maintained at 37°C in a 5% CO_2_ incubator.

### Transwell assay

Mammalian cells were seeded in 12-well plates at 2.0×10^5^ cells per well in co-culture medium for 48 h to form a confluent monolayer. Overnight shaking cultures of the indicated strains were diluted 1:100 in YLB. A volume of 0.8 mL (8.0×10^6^ CFU) of the diluted bacterial suspension was placed into the upper chamber of a transwell insert containing a 5.0 μm pore-size polyethylene terephthalate (PET) membrane (LabSelect, Beijing, China)^24^. Where indicated, 5 µg/mL of a peptide antagonist or Suramin Sodium was added simultaneously with the bacteria. The lower chamber contained sterile co-culture medium alone (no-cell control) to assess baseline bacterial migration. After 2 h of static incubation at 37 °C with 5% CO^2^, the contents of the lower chamber were collected and plated on YLB agar. CFUs were counted following 24 h incubation at 30 °C. Final CFU counts were corrected by subtracting the number of bacteria that migrated into cell-free control wells.

### Mammalian cell–bacteria co-culture and AI-2 mimic induction

Mammalian cells were seeded in 12-well plates (Sigma, Shanghai, China) at 2.0×10^5^ cells per well in 2 ml glucose-free DMEM supplemented with 20% FBS (co-culture medium) and incubated for 48 h^24^. Overnight shaking cultures of *Yptb* Δ*luxS*, *E. coli* Δ*luxS*, *V. harveyi* Δ*luxS*, *S*. Typhimurium Δ*luxS*, *P. aeruginosa,* or *K. pneumoniae* were diluted 1:100 (1.0×10^7^ CFU/mL) in YLB broth. AI-2 mimic production was induced using three distinct protocols: (1) Direct contact co-culture. Bacterial suspension (1 mL) was added directly to each well of mammalian cells. (2) Non- contact co-culture. To physically separate bacteria from mammalian cells while permitting the exchange of soluble factors, the diluted bacterial suspension was added to the upper chamber of a Transwell insert, and the insert was placed into the well containing mammalian cells. (3) Cell-free supernatant stimulation. Filter- sterilized bacterial culture supernatant (10 μL) was added directly to the mammalian cell culture. Two controls were included in parallel for all protocols: mammalian cells incubated with an equivalent volume of sterile YLB medium (referred to as “cell culture suspension”), and bacteria incubated in co-culture medium without mammalian cells. Following stimulation, all cultures were incubated at 37 °C with 5% CO₂ for 5 h. Culture supernatants were then collected and sterilized by filtration through a 0.22-µm membrane (Millipore). AI-2 mimic activity was quantified by bioluminescence reporter assay.

### AI-2 mimic production from PBS-treated cells

RAW 264.7 cells were detached from tissue culture plates with trypsin-EDTA, washed in culture medium to remove residual trypsin and serum, and then seeded in 12-well plates at 2.0×10^5^ cells per well in Dulbecco’s PBS (DPBS; Thermo Fisher Scientific)^24^. After 48 h of incubation at 37 °C and 5% CO^2^, culture supernatants were collected, sterilized through a 0.22-µm membrane (Millipore), and assayed for AI-2 mimic activity using a bioluminescence reporter assay.

### Chemotaxis assays

*Yptb* strains were grown overnight in YLB medium with shaking to an OD_600_ of 1.6, washed with M9, and resuspended in M9 medium at 1×10^8^ CFU/mL. Subsequently, 180 μL aliquots of the bacterial suspension were transferred to a 96-well plate.

Capillaries (1 μL volume, Sigma, Shanghai, China) were heat-sealed at one end and filled with cell medium, RAW 264.7 cell culture supernatant, co-culture supernatant, DPD/AI-2 (Om Scientific), mesifurane (Sigma, Cat# W366412), or PBS, then immersed in the bacterial suspension with the open ends facing downward. After 1 h at room temperature, capillaries were removed, rinsed briefly in M9, the sealed ends were broken, and the contents were expelled into 0.5 mL M9. Serial dilutions were plated on YLB agar, and the number of cells was corrected by subtracting the number of cells that entered the cell culture supernatant or PBS capillaries.

### *V. harveyi* MM32 bioluminescence reporter assay

Overnight cultures of *V. harveyi* MM32 (*luxN^−^*, *luxS^−^*) grown in AB medium were diluted 1:5,000 into AB medium, and 90 μL aliquots were dispensed into black, clear- bottom 96-well plates (Corning cat# 3603)^21^. Test solutions (10 μL per well) or buffer controls were added, and plates were incubated at 30°C with orbital shaking at 170 rpm for 10 h. Bioluminescence (counts per second) was measured using a microplate reader VictorX3 (PerkinElmer), and AI-2/AI-2 mimic activity is reported as fold induction relative to the light production induced by the buffer control.

### Quantification of AI-2 and host-derived mesifurane

To monitor AI-2 production during bacterial growth, WT *Yptb* and the Δ*luxS* mutant were cultured in YLB medium at 26 °C with shaking. At 2 h intervals, OD_600_ was recorded and culture aliquots were collected until the strains reached the stationary phase. Supernatants were sterilized by filtration and assayed for AI-2 activity using the bioluminescence reporter assay. AI-2 concentrations were determined by comparison to a known concentration of DPD/AI-2. For the host-derived mesifurane, the factor was generated according to the mammalian cell-bacteria co-culture assay. Mesifurane concentration was quantified using the same reporter assay, with concentrations determined relative to a known concentration of mesifurane.

### AI-2/ mesifurane binding assay

Protein-ligand capture was assessed by incubating DPD/AI-2, mesifurane, or co- culture supernatant with purified proteins, followed by ultrafiltration and release of bound ligand^21,24^. 3 mL of co-culture supernatant was incubated with 6 nmol of protein or BSA (negative control) overnight at room temperature with gentle agitation. Mesifurane or DPD/AI-2 was incubated with protein at an approximate 1:1 molar ratio. Following incubation, samples were concentrated using 10 kDa molecular-weight cutoff centrifugal filters (Millipore) at 4 °C and washed with PBS to remove unbound ligand. Retentates were recovered, heated at 70 °C for 5 min to denature proteins, and briefly centrifuged to pellet precipitated protein. The resulting supernatants were tested for AI-2 or mesifurane activity using the *V. harveyi* MM32 bioluminescence assay.

### Isothermal titration calorimetry (ITC)

ITC experiments were performed at 20°C using a Nano ITC Standard Volume isothermal calorimeter (TA Instruments, New Castle, DE)^21^. All proteins were dialyzed into ITC buffer (50 mM Tris, 150 mM NaCl, 10% glycerol, pH 8.0). Ligand solutions were prepared in the same buffer. The sample cell was loaded with 10 µM protein (1 mL), and the syringe was filled with co-culture supernatant, 100 µM mesifurane, 100 µM DPD/AI-2, 100 µM peptide antagonist or 100 µM Suramin Sodium (250 µL). For all experiments, the stirring speed was 200 rpm. A total of 25 injections were performed per run. Control titrations of each ligand or supernatant into buffer alone were performed to measure heats of dilution. Data were analyzed using NanoAnalyze software version 3.4, fitting the data to a one-site independent binding model. The heat of dilution was corrected by subtracting the dilution heats prior to fitting.

### Cell adhesion assay

For the quantitative adhesion assay, mammalian cells were seeded in 12-well plates at 1×10^5^ cells per well^72^. The monolayers were washed with PBS and maintained in antibiotic-free DMEM. Overnight shaking cultures of *Yptb* were washed, resuspended in PBS, and added to cells at an MOI of 100:1. Following 1 h incubation at 37 °C, non-adherent bacteria were removed by PBS washes. Cells were lysed with 1 mL of 0.1% Triton X-100 for 5-10 min at room temperature. Lysates were plated on YLB agar and enumerated for CFUs.

For the imaging-based adhesion assay, mammalian cells were seeded at 2×10^5^ cells per well in glass-bottom confocal culture dishes (Biosharp). Monolayers were infected at an MOI of 100 with *Yptb* strains expressing GFP from plasmid pKT100- GFP. Following 30 min at 37 °C, CellTracker™ Deep Red (Thermo Fisher Scientific) was added to label host cells according to the manufacturer’s instructions, and incubation continued for 30 min. Monolayers were washed thoroughly with PBS and immediately imaged on a spinning disk confocal microscope (Andor Revolution XD).

### Molecular docking analysis

The solution structure of the domain of *Yptb* YcrA, NarX and CusR were retrieved from the AlphaFoldDB under the accession code A0A0H3B5X1 (YcrA, https://alphafold.ebi.ac.uk/entry/A0A0H3B5X1),-A0A0H3B497-(NarX, https://alphafold.ebi.ac.uk/entry/A0A0H3B497),-and A0A0H3B3I9-(CusR, https://alphafold.ebi.ac.uk/entry/A0A0H3B3I9). Potential pockets and cavities were predicted using the web-based POCASA 1.1 with a probe radius of 2 Å^73^. The structure of mesifurane was downloaded from the ZINC database (ZINC1850844) and its flexible torsions were assigned using Autodock 4.42^74^. Docking simulation was done by using AutoDock Vina 1.2.0^75^, with the best binding mode selected based on the lowest docking energy. The three-dimensional figure was displayed with PyMOL v2.5.2 (http://www.pymol.org).

### Peptide antagonist design and drug screening

For peptide antagonist design, a 3D model of YcrA was constructed, and residues 116-144 and 165-170 were defined as the binding region, with A120, A122, A138, A140, and A167 assigned as hotspot residues for mesifurane interaction. RFdiffusion was used to generate peptides that occlude this active-site region^76^. The designed peptides were docked with Hdock, and the top 30 by docking score were retained^77^. These candidates were evaluated by one-to-one complex prediction with YcrA using AlphaFold 3, ranked by combined pTM and ipTM scores^78^, and a final peptide antagonist library was established.

For drug screening, high-resolution structures of YcrA orthologs from *A. baumannii*, *E. coli*, and *P. aeruginosa* were retrieved from the PDB database or predicted with AlphaFold 3^79^. Homologous active-site pockets involved in mesifurane binding were identified by structural comparison, marked, and saved in PDBQT format. A library of ∼26,000 clinical and development-stage compounds underwent standardized preprocessing and was likewise converted to PDBQT format. High-throughput virtual screening was performed by semi-flexible docking with AutoDock Vina 1.2.0^75^ against the three ortholog pockets, yielding a high-affinity compound library. Candidate compounds were then individually modeled in complex with each protein using AlphaFold 3,^78^ and the occlusion effect and binding affinity were evaluated under strict spatial constraints on the active pockets, ultimately defining a drug library of molecules that simultaneously display high affinity for all three orthologs.

### HPLC analysis

Filtered supernatants were injected into an HPLC (LC-30A, Shimadzu) system equipped with a C18 reversed-phase column and a UV detector. Components were eluted isocratically with 55% A (methanol) and 45% B (0.1% phosphoric acid aqueous solution) in 8 min at a flow rate of 1 mL/min^80^. The detection wavelength was 280 nm. Mesifurane was run as standard. The levels of synthesized mesifurane in the supernatants were determined from the standard curve obtained using known concentrations of mesifurane.

### LC-MS analysis

The filtered supernatants and mesifurane standard were analyzed using a Liquid chromatography-quadrupole ion trap-mass spectrometry (LC-QTRAP-MS, QTRAP5500, AB SCIEX) system to quantify mesifurane production by the cells and to evaluate and compare the synthesis levels of this metabolite under different treatment conditions.

### RNA-seq analysis

*Yptb* WT strains were grown to the exponential phase in YLB medium and treated with or without host-bacteria co-culture supernatant for 2 h. Bacterial cells were harvested and then total RNA was extracted using the RNeasy mini kit (Qiagen). Genomic DNA was degraded using the TURBO DNA-free™ kit (Life Technologies). Subsequently, ribosomal RNA was removed using MICROBExpress™ Bacterial mRNA Enrichment Kit (Life Technologies). cDNA libraries were prepared using the Bacterial ScriptSeq Complete Kit (Illumina) following the manufacturer’s instructions, and sequencing was performed by Sangon Biotech Co., Ltd. (Shanghai, China). RNA-seq was performed on triplicate samples. Sequence reads were mapped to the *Y. pseudotuberculosis* YPIII reference genome NZ_CP009792.1 [https://www.ncbi.nlm.nih.gov/nuccore/NZ_CP009792.1] using Bowtie (version 2.3.2). FeatureCounts v1.6.0 was used to count the number of reads mapped to each gene. Relative transcript abundance was calculated based on the number of reads per kilobase per million mapped sequence reads (RPKM). Differential expression analysis was performed using the DESeq2 R package (version 1.12.4), with a threshold set to p < 0.05 and |log2 foldchange| > 1. P values were adjusted using the Benjamini–Hochberg approach to control the false discovery rate (FDR)^81^. The role of the differentially expressed genes was investigated using Kyoto Encyclopedia of Genes and Genomes (KEGG) pathway analysis, considering pathways with P-values below 0.05 as significantly enriched. The raw FASTQ files for the RNAseq libraries have been deposited in the NCBI Sequence Read Archive (SRA) under the BioProject accession number PRJNA1359123.

### Bacterial RNA extraction and qRT-PCR analysis

Cultures of *Yptb* strains were harvested, and then total RNA was extracted by RNAprep Pure Cell/Bacteria Kit (Tiangen Biotech). After treatment with RNase-free DNase I (Sigma Aldrich), cDNA was reverse transcribed from RNA by TransScript II One-Step gDNA Removal and cDNA Synthesis SuperMix (TransGen Biotech). qRT- PCR was performed, and the relative abundance of 16S rRNA was used as an internal standard.

### T6SS activity analysis

To quantify T6SS transcriptional activation during infection, RAW 264.7 cells were seeded in 12-well plates at 1×10^6^ cells per well. Overnight cultures of *Yptb* were washed and added to cells at an MOI of 100:1 for 2 h at 37°C, 5% CO^2^. After infection, the entire well content (supernatant and cell layer) was collected. Cells were lysed with 0.1% Triton X-100 to release cell-associated bacteria. Bacterial cells were pelleted by centrifugation, and T6SS gene expression was quantified by qRT- PCR.

For fluorescent reporter visualization of T6SS activity, a translational fusion of the T6SS ATPase gene *clpV4* to superfolder GFP (sfGFP) on a plasmid was constructed^39,82^. The reporter plasmid was introduced into *Yptb* strains by conjugation via *E. coli* strain S17-1 λpir. For imaging, cells were seeded in glass-bottom confocal dishes at 1 × 10^5^ cells per well. Monolayers were infected at an MOI of 100 with mid-exponential phase (OD_600_ ≈ 0.8) reporter *Yptb* strains. Following 90 min at 37 °C, CellTracker™ Deep Red was added to label host cells, and incubation continued for 30 min. Monolayers were washed extensively with PBS and immediately imaged on a spinning-disk confocal microscope.

### Cell apoptosis assay

Apoptosis was quantified using an Annexin V-FITC/PI Apoptosis Detection Kit (Beyotime Biotechnology, #C1062M). Mammalian cells (1×10^6^) were infected with the indicated strains at MOI 100 for 2 h at 37°C. Where indicated, 5 µg/ml of a peptide antagonist or Suramin Sodium was added either to the cells or concurrently with bacterial inoculation. Cells were harvested, washed with PBS, resuspended in 195 μL binding buffer, and stained with Annexin V-FITC and PI for 10-20 min at room temperature in the dark. Samples were analyzed immediately on a CytoFLEX flow cytometer (Beckman Coulter), and the data were processed using CytExpert software.

For microscopic visualization of apoptosis, mammalian cells were seeded in glass-bottom confocal dishes at 2×10^5^ cells per well. Cells were infected at an MOI of 100 with *Yptb* strains expressing GFP from plasmid pKT100-GFP. Following 90 min at 37 °C, CellTracker™ Deep Red was added to label host cells, and incubation continued for 30 min. Monolayers were washed, stained with PI for 10 min at room temperature, and immediately imaged using a spinning-disk confocal microscope.

### Chromosomal fusion strains construction and β-galactosidase assay

The promoter regions of T6SS-4 and *luxS* were amplified by PCR and separately cloned into the *lacZ* fusion reporter vector pDM4-*lacZ*. Recombinant plasmids were transformed into *E. coli* strain S17-1 λpir and then transferred to *Yptb* strains by conjugation. The transconjugants were selected on YLB agar containing nalidixic acid and chloramphenicol. The *lacZ* fusion reporter strains were grown to the exponential phase in M9 medium and treated with or without co-culture supernatant. β-galactosidase activity was assayed using ONPG (o-nitrophenyl-β- dgalactopyranoside) as the substrate according to the Miller method^83^.

### Western blot analysis

Samples were separated by sodium dodecyl sulfate-polyacrylamide gel electrophoresis (SDS-PAGE) and transferred onto polyvinylidene fluoride (PVDF) membranes (Millipore). The membrane was blocked with QuickBlock Blocking Buffer (Beyotime) for 30 min at room temperature and incubated with primary antibodies at 4°C overnight. The membrane was washed three times in TBST buffer with Tween 20 (50 mM Tris, 150 mM NaCl, and 0.05% Tween 20, pH 7.4) and incubated with horseradish peroxidase-conjugated secondary antibodies for 1 h. Signals were detected using the ECL Kit (Invitrogen) following the manufacturer’s specified protocol.

### In vitro phosphorylation assay

To generate phosphorylated NarP (NarP-P), 4 μM NarP was incubated with acetyl phosphate at a molar ratio of 1:1200 in phosphorylation buffer (40 mM KCl, 50 mM Tris, pH 7.5, 40 mM MgCl^2^, 2.5% Glycerol) for 20 min^84^. For kinase assays, reactions contained 4 µM NarX, 2 µM NarP, 4 µM ATP, and co-culture supernatant (or cell-culture supernatant as a control) in the same phosphorylation buffer. Reactions were incubated for 30 min to permit NarP phosphorylation. Negative controls omitted either ATP or NarX. Following incubation, all phosphorylation reactions were terminated, and the phosphorylation levels were quantified by Ultra-High Resolution Liquid Chromatography-Mass Spectrometry (LC-MS) using a Thermo Fisher Fusion Lumos system.

### Phos-tag SDS-PAGE analysis

Phos-tag SDS-PAGE analysis was conducted using the user manual for Phos-tag Acrylamide AAL-107 (Wako Pure Chemical Industries, Osaka, Japan) with minor modifications^85^. *Yptb* strains harboring pKT100-*narP*-*his^6^* were grown to the exponential phase in M9 medium and treated with co-culture supernatant, 10 μM mesifurane, or vehicle control for 2 h at room temperature. Total proteins were extracted from the cell pellets and analyzed using both conventional and Phos-tag SDS-PAGE. Phos-tag gels consisted of a 5% acrylamide stacking gel and an 8% acrylamide separating gel containing 50 µM Phos-tag ligand and 100 µM MnCl^2^. Following electrophoresis, gels were washed three times in transfer buffer containing EDTA to chelate Mn^2+^, then once in EDTA-free buffer.

Immunoblot analysis was performed to detect the phosphorylation levels of PilG using an anti-His antibody.

### ChIP-qPCR

*Yptb* strains expressing in situ tagged FLAG-NarP and FLAG-CusR were generated by using the CRISPR-Cas9 system^86^. After incubation with co-culture supernatant for 2 h, cultures were treated with 1% formaldehyde for 15 min at room temperature. The crosslinking reaction was stopped by 125 mM glycine. Cells were washed with PBS, resuspended in lysis buffer (25 mM Tris-HCl, pH 7.5; 150 mM NaCl; 1 mM EDTA; 0.1% Triton X-100; 0.1% SDS), and sonicated to generate DNA fragments of 100- 500 bp. Cell debris was removed by centrifugation at 4°C, and the supernatant was used as the input sample for the IP experiments. Subsequently, the input sample was incubated with either no antibody (mock-IP) or mouse anti-FLAG (1:1,000 dilution, Abways, Shanghai, China, cat# AB0028), followed by enrichment with Protein A magnetic beads (Sigma-Aldrich). Immunoprecipitated Protein-DNA complexes were eluted from the beads using elution buffer (50 mM Tris-HCl, pH 7.5; 10 mM EDTA; 1% SDS) at 65°C for 30 min. Crosslinks were reversed by incubation for 6 h at 65°C in 0.5×elution buffer plus 250 μg/ml proteinase K. DNA was extracted twice with phenol/chloroform (1:1), precipitated and resuspended in distilled water^87^. Enriched DNA was quantified by qPCR using KAPA SYBR FAST qPCR Kit (Kapa Biosystems) in a Roche LightCycler 96 Real-Time PCR system. Signals were normalized to input DNA^88^, and background from mock-IP samples was subtracted.

### EMSAs

DNA probes were amplified by PCR and purified on 6% native polyacrylamide gels. In a 20 μl reaction system, 50 ng or 20 ng of DNA probes were mixed with the indicated proteins in a Tris buffer (20 mM TrisHCl, 4 mM MgCl^2^, 100 mM NaCl, 1 mM dithiothreitol, 1% NP-40, and 10% glycerol, pH 7.4). Where indicated, ATP, co- culture supernatant, or mesifurane were added simultaneously with the indicated proteins to the reaction system. After incubation for 30 min at room temperature, the reaction mixtures were subjected to 6% native polyacrylamide gel and run in 0.5× TBE buffer at 100 V. The DNA probe was detected using SYBR Safe DNA gel stain (Invitrogen) and imaged by the Tanon gel analysis software version 2.30 (Tanon 5200Multi, China).

### Bacterial growth curves

Growth kinetics of *Yptb* WT and Δ*ycrA* were monitored in YLB medium at 26 °C. *E. coli*, *P. aeruginosa*, and *A. baumannii* were grown in LB medium at 37 °C with the indicated concentrations of Suramin Sodium. Overnight cultures of all strains were diluted 1:100 in the appropriate fresh medium, and 200 μL was dispensed into 96- well plates in triplicate. Plates were incubated with continuous shaking in a microplate reader, and OD_600_ was measured every 2 h.

### Mouse infection

All animal care and experimental procedures were performed in accordance with the Regulations for the Administration of Affairs Concerning Experimental Animals approved by the State Council of the People’s Republic of China. The protocol was approved by the Animal Welfare and Research Ethics Committee of Northwest A&F University (Protocol number: XN2023-1004). Female BALB/c mice (5-6 weeks old) were housed under a 12 h light/dark cycle at 24 ± 2 °C and 50–60% humidity with a 3-day acclimation period prior to experiments. *Yptb* was grown to mid-exponential phase in YLB medium at 26 °C. *E. coli*, *A. baumannii* and *P. aeruginosa* were grown in LB at 37°C. Strains were washed with PBS and used for infection.

For survival assays. (1) Intragastric infection: Mice (n = 12 per group) were inoculated with 1 × 10^9^ CFU of the indicated *Yptb* strains (100 μL) by oral gavage. Survival was monitored twice daily for 21 days^89^. (2) Intraperitoneal infection: Mice (n = 10 per group) were injected intraperitoneally with 1 × 10^7^ CFU of the indicated *Yptb* strains (100 μL). Survival was recorded twice daily for 6 days. (3) Peptide antagonist therapeutic intervention: For each bacterial pathogen, mice were randomly allocated to four groups (n = 10 per group): (i) infection + PBS, (ii) infection + peptide antagonist, (iii) uninfected + PBS, and (iv) uninfected + peptide antagonist. Infection was performed via intraperitoneal injection of 100 μL *Yptb* (1 × 10^7^ CFU), *E. coli* (1 × 10^7^ CFU), or *A. baumannii* (5 × 10^5^ CFU). Concomitantly, mice received an intraperitoneal injection of either peptide antagonist (10 mg/kg) or an equivalent volume of PBS (100 μL). This treatment was repeated every 24 h for 6 days. Survival was recorded twice daily throughout the 6-day observation period^90^. (4) Suramin Sodium therapeutic intervention: For each bacterial pathogen, mice were randomly allocated to four groups (n = 10 per group): (i) infection + PBS, (ii) infection + Suramin Sodium, (iii) uninfected + Suramin Sodium, and (iv) uninfected + PBS. Infection was performed via intraperitoneal injection of 100 μL *E. coli* (2 × 10^7^ CFU), *P. aeruginosa* (5× 10^6^ CFU), or *A. baumannii* (1× 10^6^ CFU). At the designated time point (2 h or 8 h) post-infection, mice received a subcutaneous injection of Suramin Sodium (10 mg/kg) or an equivalent volume of PBS (100 μL). This treatment was administered every 12 h for the first 3 days, after which the dosing frequency was shifted to every 24 h. Survival was monitored twice daily for 6 days.

Bacterial Load quantification. (1) Intragastric infection: Mice were inoculated with 1 × 10^9^ CFU of the indicated *Yptb* strains (100 μL) by oral gavage (n = 8 per group). For fecal load, feces were collected from live mice at 48 h post-infection, weighed, and homogenized in PBS. For tissue load, mice were euthanized by CO^2^ asphyxiation followed by cervical dislocation at 48 h. The cecum and small intestine were weighed and homogenized in PBS, and serial dilutions of the homogenates were plated on YLB plates with 20 μg/mL nalidixic acid. CFU were counted and are given as CFU per gram of organ/tissue. (2) Bacterial burden following therapeutic intervention: Mice were infected and treated as described in Suramin Sodium therapeutic intervention, with the following groups (n = 6 per group): infection + PBS and infection + Suramin Sodium. At 12 h post-treatment (14 h post-infection), blood was collected, and mice were euthanized. Peritoneal lavage was performed with 5 mL of sterile PBS, and the lavage fluid was harvested. Spleen and liver were collected, weighed, and homogenized in PBS. Serial dilutions of blood, lavage fluid, and tissue homogenates were plated on LB plates. CFU were enumerated and expressed as CFU per mL for blood and lavage fluid, or as CFU per gram of tissue for spleen and liver.

### ELISA quantification of cytokines

Mice were infected and treated as described in Suramin Sodium therapeutic intervention, with the following groups (n = 6 per group): uninfected control (PBS only), infection + PBS, and infection + Suramin Sodium (initiated at 2 h post-infection). At 12 h post-treatment (14 h post-infection), blood was collected and allowed to clot at room temperature for 30 min before centrifugation (2,000 × g, 10 min, 4 °C) to obtain serum. Peritoneal lavage was performed with 5 mL of sterile PBS, and the lavage fluid was centrifuged to remove cellular debris. Spleen and liver were harvested, weighed, and homogenized in Cell Lysis Buffer for Western and IP (Beyotime) supplemented with protease inhibitors^91,92^. Tissue homogenates were centrifuged (12,000 × g, 10 min, 4 °C), and the supernatants were collected. Levels of IL-1β, IL-6, IL-10, TNF-α, and IFN-γ in serum, peritoneal lavage fluid, and tissue lysates were measured using commercially available mouse ELISA kits (Beyotime) according to the manufacturer’s instructions^93^. Absorbance was read at 450 nm on a microplate reader. Cytokine concentrations were calculated from standard curves and normalized to the original sample volume (blood, peritoneal lavage fluid, tissue homogenate). For liver and spleen, homogenates were prepared at 0.6 g/mL and 0.12 g/mL, respectively.

### Tissue and blood RNA extraction and qRT-PCR analysis

Mice were infected, treated, and sacrificed as described in the ELISA quantification of cytokines. Total RNA from blood was extracted using the RNAeasy™ Blood RNA Isolation Kit with Spin Column (Beyotime). Peritoneal lavage was performed with 5 mL of sterile PBS, and the lavage fluid was centrifuged to collect peritoneal cells. Peritoneal cells, spleen, and liver were processed, and total RNA was extracted using the RNAeasy™ Animal RNA Isolation Kit with Spin Column (Beyotime)^94^. RNA was reverse-transcribed into cDNA using the TransScript II One-Step gDNA Removal and cDNA Synthesis SuperMix (TransGen Biotech)^95^. qRT-PCR was performed, and the relative expression of *Actb* (encoding β-actin) was used as the internal control for normalization.

### Evaluation of Bacterial Resistance Development

*E. coli, P. aeruginosa,* and *A. baumannii* were serially passaged for approximately 200 generations^96^ in the absence (serial passage) or presence (serial passage under drug pressure) of 10 μg/mL suramin sodium. The ancestral strain was maintained without passage.

The in vivo efficacy of suramin sodium against the ancestral strain and both passaged populations was evaluated using the mouse infection model described above. Briefly, for each bacterial pathogen and each population, mice were randomly allocated to the following groups (n = 10 per group): infection + PBS, infection + suramin sodium (initiated at 2 h post-infection). Survival was monitored for 6 days.

### Statistical analysis

GraphPad Prism Software (GraphPad Prism 8.00) or Microsoft Excel 2021 was used to perform all statistical analyses. For ITC experiments, standard deviation was calculated from replicate experiments. Data from the Mouse Infection assay were analyzed using the Log-rank (Mantel–Cox) test. All other experiments were analyzed using the two-tailed unpaired Student’s *t* test. Data are presented as mean ± SD and statistical significance was defined by *P* < 0.05 (*), *P* < 0.01 (**), *P* < 0.001 (***), and *P* < 0.0001 (****).

### Data availability

The protein 3D coordinate data used in this study are available in the AlphaFoldDB database under accession codes A0A0H3B5X1,A0A0H3B497, and A0A0H3B3I9. The RNA-seq data generated in this study have been deposited in the NCBI BioProject database under the accession number PRJNA1359123. Other data supporting the findings of this study are provided within the article and its Supplementary Information files, or can be obtained from the corresponding authors upon request.

## Acknowledgments

This work was supported by grants from the National Natural Science Foundation of China (Grant 32330004 to X.S., and Grant 32300038 to S.L.). The Shaanxi Fundamental Science Research Project for Chemistry & Biology (Grant No. 22JHZ008 to X.S.), the China Postdoctoral Science Foundation (Grant 2026M792986 to Q.L.), Ningbo Top Medical and Health Research Program (No.2023020713 to X.S. and Y.C.), and Science and Technology Innovation Yongjiang 2035 Key Research and Development Project of Ningbo (No. 2025Z180 to X.S. and Y.C.). We thank the Teaching and Research Core Facility at College of Life Science (Xiyan Chen, Ningjuan Fan, and Hui Duan), Life Science Research Core Services (Luqi Li and Min Zhou), Plant Protection Science and Technology Innovation Platform (Zhimei Bai and Yan Li), and State Key Laboratory for Crop Stress Resistance and High-Efficiency Production (Hua Zhao), Northwest A&F University for technical support. We also thank Beyotime Biotech Inc for helping in CRISPR-Cas9 genome editing in RAW 264.7 cells.

## Author contributions

Author contributions: X.S., Q.L., S.L. designed research; Q.L., S.L., Y.Z., S.C., H.S., Y.Y., T.H., Z.Wei., Z.Wang., Y.W., L.Y. and C.Y. performed research; Q.L., S.L., and Y.Z. analyzed data; and X.S., Q.L., and S.L. wrote the paper.

## Declaration of interests

Patent applications covering aspects of this work are currently pending. The authors declare no other competing interests.

## Notes

### Competing Interest Statement

The authors have declared no competing interest.

### Summary of Updates

Author list updated; Funding Information updated; Figure 1, 3 revised; Figure 7 added; Supplemental files updated; The writing of the entire manuscript, including the title, has been revised and improved.

