## Supplementary Information for "Bacteria Hijack a Host Metabolite via Three Distinct Sensors to Orchestrate Infection"

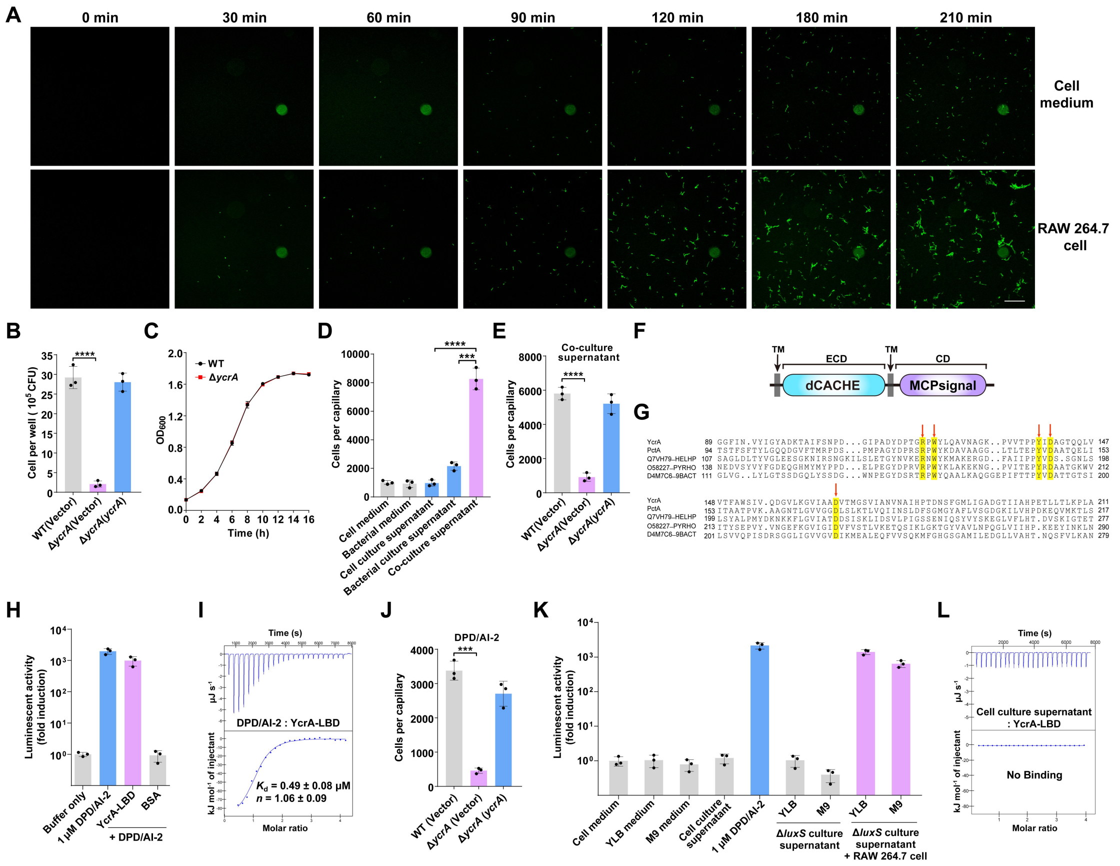


**Figure S1. YcrA mediates *Yptb* chemotaxis toward macrophages and AI-2.**

**(A)** Time-lapse visualization of *Yptb* migration toward macrophages in a transwell system. GFP-tagged *Yptb* was added to the upper chamber, while a RAW 264.7 monolayer was established in the lower well. Images were acquired every 30 min. Sterile cell medium (cell-free) served as a migration control. Images are representative of three independent experiments with similar results. Scale bar, 50 μm.

**(B)** YcrA is essential for chemotaxis toward macrophages. Transwell assays comparing migration of the indicated strains toward RAW 264.7 cells after 2 h. Migrated bacteria were quantified, and final CFU counts were corrected by subtracting the number of bacteria that migrated into cell-free control wells.

**(C)** Growth curves of *Yptb* WT and Δ*ycrA* were monitored in YLB medium at 26 °C,180rpm. $OD_{600}$ was measured every 2 h.

**(D)** Chemotactic response of WT *Yptb* to cell medium, bacterial medium, cell culture supernatant, bacterial culture supernatant, or host-bacterial co-culture supernatant, assessed by quantitative capillary assay. The data represent bacterial accumulation in capillaries.

**(E)** YcrA mediates chemotaxis toward host-bacteria co-culture supernatant. Quantitative capillary assays measured the accumulation of indicated *Yptb* strains in capillaries filled with co-culture supernatant for 1 h. The final CFU counts were corrected by subtracting the number of cells that entered the cell culture supernatant capillaries.

**(F)** Schematic domain organization of YcrA. TM transmembrane domain, ECD extracytoplasmic domain, CD cytoplasmic domain.

**(G)** Multiple sequence alignment of dCache ligand-binding domains (LBDs) from YcrA and homologous proteins. Highly conserved residues R120, W122, Y138, D140 and D167 are highlighted in yellow. Alignment was performed using ClustalW and visualized with GeneDoc 2.7.

**(H)** YcrA captures bacterial AI-2. Purified YcrA-LBD was incubated with DPD/AI-2; the captured ligand was heat-eluted and quantified via the *V. harveyi* MM32 bioluminescence reporter assay. 1 μM DPD/AI-2 served as a positive control, and BSA served as a negative control. Data are fold induction relative to the buffer control.

**(I)** ITC analysis of the binding affinity between YcrA-LBD and AI-2. The displayed thermogram is one representative of three independent experiments; $K_d$ and binding stoichiometry (*n*) are presented as mean ± s.d. from three independent experiments.

**(J)** YcrA is required for AI-2-induced chemotaxis. Quantitative capillary assays measured accumulation of indicated *Yptb* strains in capillaries containing 10 μM DPD/AI-2 for 1 h. The number of cells was corrected by subtracting the number of cells that entered the PBS capillaries.

**(K)** Macrophages produce an AI-2 mimic upon stimulation by *Yptb* culture suspension. *Yptb* Δ*luxS* was grown overnight in YLB or M9*.* Filter-sterilized bacterial culture supernatant (10 μL) was added directly to RAW 264.7 cells. After 5 h of stimulation, AI-2 mimic activity in filter-sterilized supernatants was measured using the *V. harveyi* MM32 bioluminescence reporter strain. Supernatants from *Yptb* Δ*luxS* cultured alone or RAW 264.7 cells cultured alone showed no detectable activity. 1 μM DPD/AI-2 was the positive control. Data are fold induction relative to the buffer control.

**(L)** No detectable binding between YcrA and cell culture supernatant. The binding affinity was determined using ITC. The displayed thermogram is one representative of three independent experiments.

Data are represented as mean ± SD of three biological replicates, each with three technical replicates. Statistical significance was determined with the two-tailed unpaired Student's *t*-test. ***$P < 0.001$; ****$P < 0.0001$.

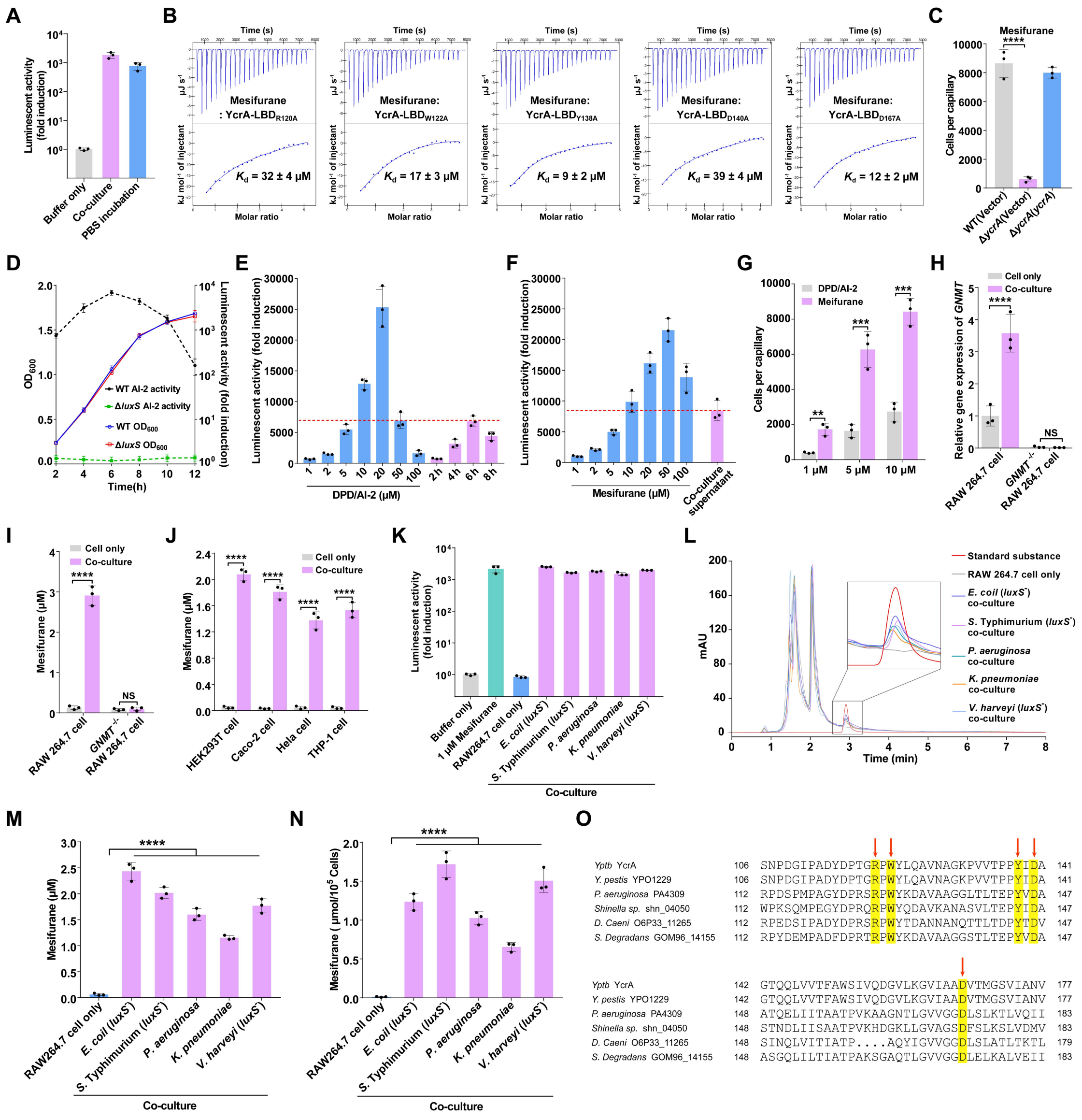


**Figure S2. YcrA homologs capable of sensing the mesifurane are widespread in bacteria.**

**(A)** RAW 264.7 cells produce mesifurane in PBS. RAW 264.7 cells were cultured in PBS for 48 h. Culture supernatants were collected, and mesifurane activity was quantified using the *V. harveyi* MM32 bioluminescence reporter assay. Co-culture supernatant served as a positive control. Data are fold induction relative to the buffer control.

**(B)** ITC analysis of the binding affinity between YcrA-LBDs point mutants and mesifurane. The displayed thermogram is one representative of three independent experiments; $K_d$ and binding stoichiometry ($n$) are presented as mean ± s.d. from three independent experiments.

**(C)** YcrA is essential for chemotaxis toward mesifurane. Quantitative capillary assays measured accumulation of indicated *Yptb* strains in capillaries containing 10 μM mesifurane for 1 h. The number of cells was corrected by subtracting the number of cells that entered the PBS capillaries.

**(D)** AI-2 activity during growth of WT *Yptb* and the Δ*luxS* mutant. Growth ($OD_{600}$, left axis) and AI-2 activity (right axis) in supernatants of WT *Yptb* and the Δ*luxS* mutant grown in YLB at 26°C. AI-2 activity was quantified using the *V. harveyi* MM32 bioluminescence reporter assay, and data are fold induction relative to the buffer control. Solid lines represent $OD_{600}$, while dashed lines indicate AI-2 activity.

**(E)** Determine the maximum concentration of AI-2 produced by *Yptb.* AI-2 activity generated at different time points was determined by comparison to a known concentration of DPD/AI-2 standard.

**(F)** Determine the maximum concentration of mesifurane produced by the host cell. Mesifurane activity generated by the *Yptb* Δ*luxS*-RAW 264.7 cells co-culture was determined by comparison to a known concentration of mesifurane standard.

**(G)** *Yptb* exhibits stronger chemotaxis toward mesifurane than DPD/AI-2 at equivalent concentrations. Quantitative capillary assays measured accumulation of *Yptb* in capillaries containing the indicated concentrations of DPD/AI-2 or mesifurane for 1 h. The number of cells was corrected by subtracting the number of cells that entered the PBS capillaries.

**(H)** qRT-PCR analysis of the *GNMT* mRNA levels in WT and *GNMT*$^{-/-}$ RAW 264.7 cells left untreated or infected with Δ*luxS* for 5 h. Expression was normalized to *Actb* and is presented as fold change relative to uninfected WT cells.

**(I and J)** Quantification of mesifurane by HPLC. Mesifurane concentrations in the indicated samples were determined by peak-area integration against a calibration curve generated with authentic mesifurane standards.

**(K)** Multiple bacterial species induce mesifurane production in macrophages. RAW

264.7 cells were co-cultured with *Escherichia coli* BL21 (*luxS*$^{-}$), *Salmonella* Typhimurium SL1344 (*luxS*$^{-}$), *Pseudomonas aeruginosa* PAO1, *Klebsiella pneumoniae*, and *Vibrio harveyi* MM32 (*luxS*$^{-}$); co-cultured supernatants were assayed for mesifurane activity using the *V. harveyi* MM32 reporter. 1 μM mesifurane served as a positive control, and BSA served as a negative control. Data are fold induction relative to the buffer control.

**(L)** HPLC fractionation of RAW 264.7 cell supernatants. Overlaid chromatograms show mesifurane standard (red), co-culture supernatants from RAW 264.7 cells with *Escherichia coli* (dark blue), *Salmonella* Typhimurium (purple), *Pseudomonas aeruginosa* (green), *Klebsiella pneumoniae* (orange), *Vibrio harveyi* (blue), and RAW 264.7 cell culture supernatants (grey). The active fraction is highlighted (inset). mAU, milli-absorbance units.

**(M)** Quantification of mesifurane by HPLC. Mesifurane concentrations in the indicated samples were determined by peak-area integration against a calibration curve generated with authentic mesifurane standards.

**(N)** LC-MS/MS quantification of mesifurane production. Mesifurane levels were measured in supernatants from RAW 264.7 cells cultured alone or co-cultured with the indicated bacterial strains.

**(O)** Sequence alignment of YcrA-LBD with homologs. Conserved residues R120, W122, Y138, D140, and D167 are highlighted in yellow. Alignment was performed using ClustalW and visualized with GeneDoc.

Data are represented as mean ± SD of three biological replicates, each with three technical replicates. Statistical significance was determined with the two-tailed unpaired Student's *t*-test. ****$P < 0.0001$; NS, not significant.

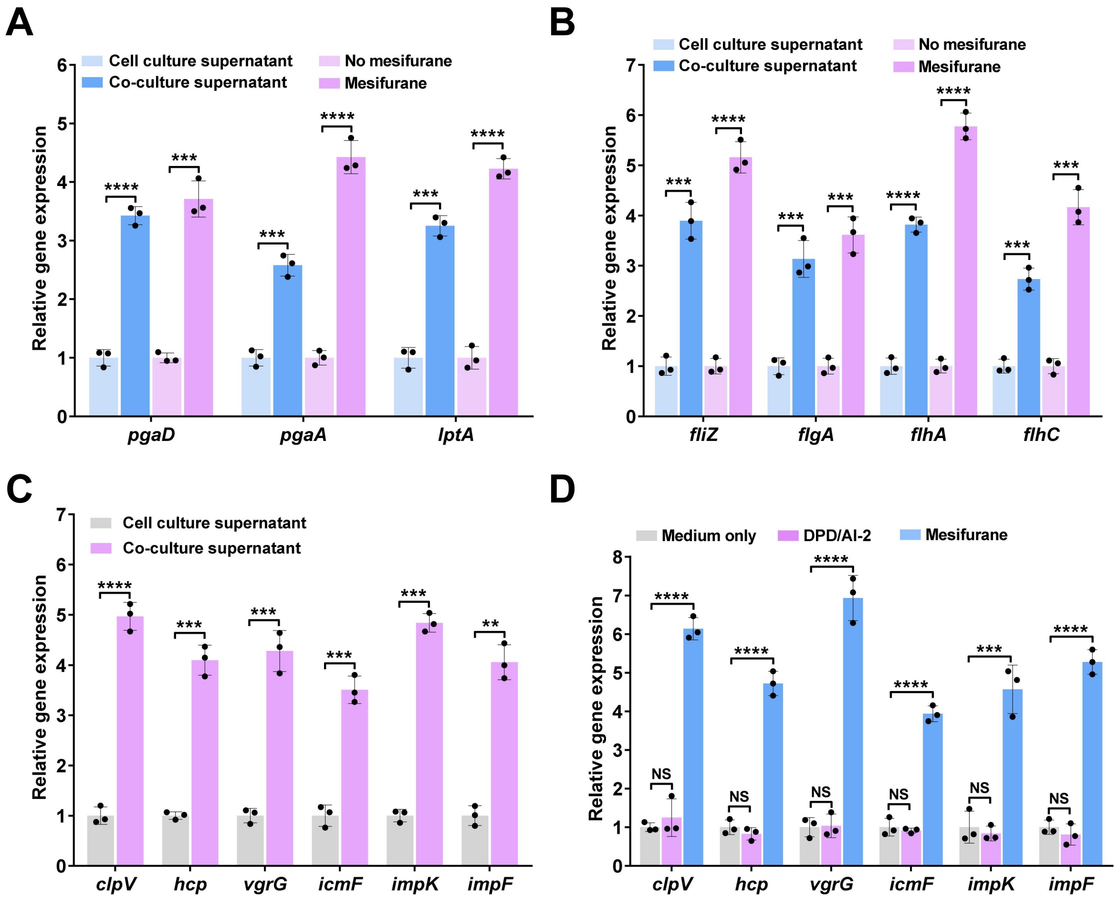


**Figure S3. Mesifurane induces the expression of biofilm, flagellar, and T6SS genes in *Yptb*.**

**(A–C)** qRT-PCR analysis of (A) biofilm-associated genes, (B) flagellar-associated genes, and (C) T6SS genes in *Yptb* after 2 h of stimulation. Cells were treated with cell culture supernatant (unstimulated control), co-culture supernatant (containing mesifurane), or 10 μM mesifurane. T6SS genes (C) were tested only with unstimulated and co-culture supernatant conditions. Expression was normalized to 16S rRNA and is presented as fold change relative to the unstimulated control.

**(D)** In the Δ*luxS* mutant, mesifurane (10 μM) strongly induced the expression of T6SS core genes, whereas DPD/AI-2 at the same concentration had no detectable effect. qRT-PCR analysis of T6SS core gene expression in *Yptb* Δ*luxS* left untreated or stimulated with 10 μM DPD/AI-2 or mesifurane for 2 h. Expression was normalized to 16S rRNA and is presented as fold change relative to the untreated control.

Data are represented as mean ± SD of three biological replicates, each with three technical replicates. Statistical significance was determined with the two-tailed unpaired Student's *t*-test. $^{**}P < 0.01$; $^{***}P < 0.001$; $^{****}P < 0.0001$; NS, not significant.

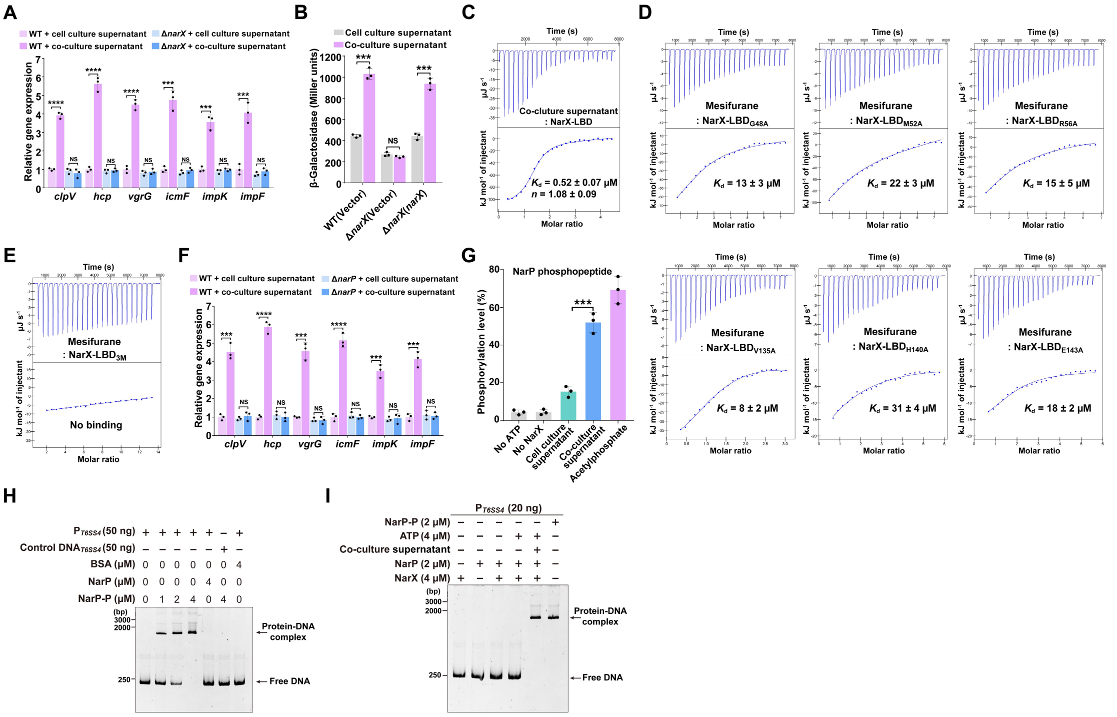


**Figure S4. The NarX-NarP two-component system mediates mesifurane to activate T6SS-4.**

**(A)** qRT-PCR analysis of T6SS genes in WT and Δ*narX* strains after 2 h of stimulation. Cells were treated with cell culture supernatant (unstimulated control) and co-culture supernatant (containing mesifurane). Expression was normalized to 16S rRNA and is presented as fold change relative to the unstimulated WT control.

**(B)** NarX is required for mesifurane-induced T6SS promoter activation. β-Galactosidase assays quantifying the activity of a chromosomal $P_{T6SS4}$*::lacZ* reporter in indicated *Yptb* strains treated with co-culture supernatant. *Yptb* strains treated with cell culture supernatant served as a control.

**(C)** ITC analysis of the binding affinity between NarX-LBD and co-culture supernatant. The displayed thermogram is one representative of three independent experiments; $K_d$ and binding stoichiometry ($n$) are presented as mean ± s.d. from three independent experiments.

**(D and E)** ITC analysis of mesifurane binding to NarX-LBD point mutants (D) and the NarX-LBD$_{3M}$ triple mutant (E). The displayed thermogram in (D, E) is one representative of three independent experiments; $K_d$ and binding stoichiometry ($n$)

are presented as mean ± s.d. from three independent experiments.

**(F)** qRT-PCR analysis of T6SS genes in WT and Δ*narP* strains after 2 h of stimulation. Cells were treated with cell culture supernatant (unstimulated control) and co-culture supernatant (containing mesifurane). Expression was normalized to 16S rRNA and is presented as fold change relative to the unstimulated WT control.

**(G)** Mesifurane stimulates NarX-dependent NarP phosphorylation. Purified NarP and NarX were incubated with ATP in the presence or absence of co-culture supernatant for 40 min. Reactions containing acetyl phosphate served as a positive control for NarP phosphorylation; reactions lacking ATP or NarX served as a negative control. Phosphorylation of NarP was quantified by LC-MS/MS analysis.

**(H)** Phosphorylated NarP binds the T6SS-4 promoter. EMSA showing binding of Acetyl phosphate-phosphorylated NarP to the T6SS-4 promoter probe. BSA and coding-region DNA fragments served as negative controls.

**(I)** NarP binds the T6SS-4 promoter in vitro. EMSA was performed with NarP and the T6SS-4 promoter probe in the presence or absence of co-culture supernatant and/or NarX. Acetyl phosphate-phosphorylated NarP was included as a positive control. The gel shown in (H and I) is representative of three independent experiments with similar results.

Data are represented as mean ±SD of three biological replicates, each with three technical replicates. Statistical significance was determined with the two-tailed unpaired Student's *t*-test. ***$P < 0.001$; ****$P < 0.0001$; NS, not significant.

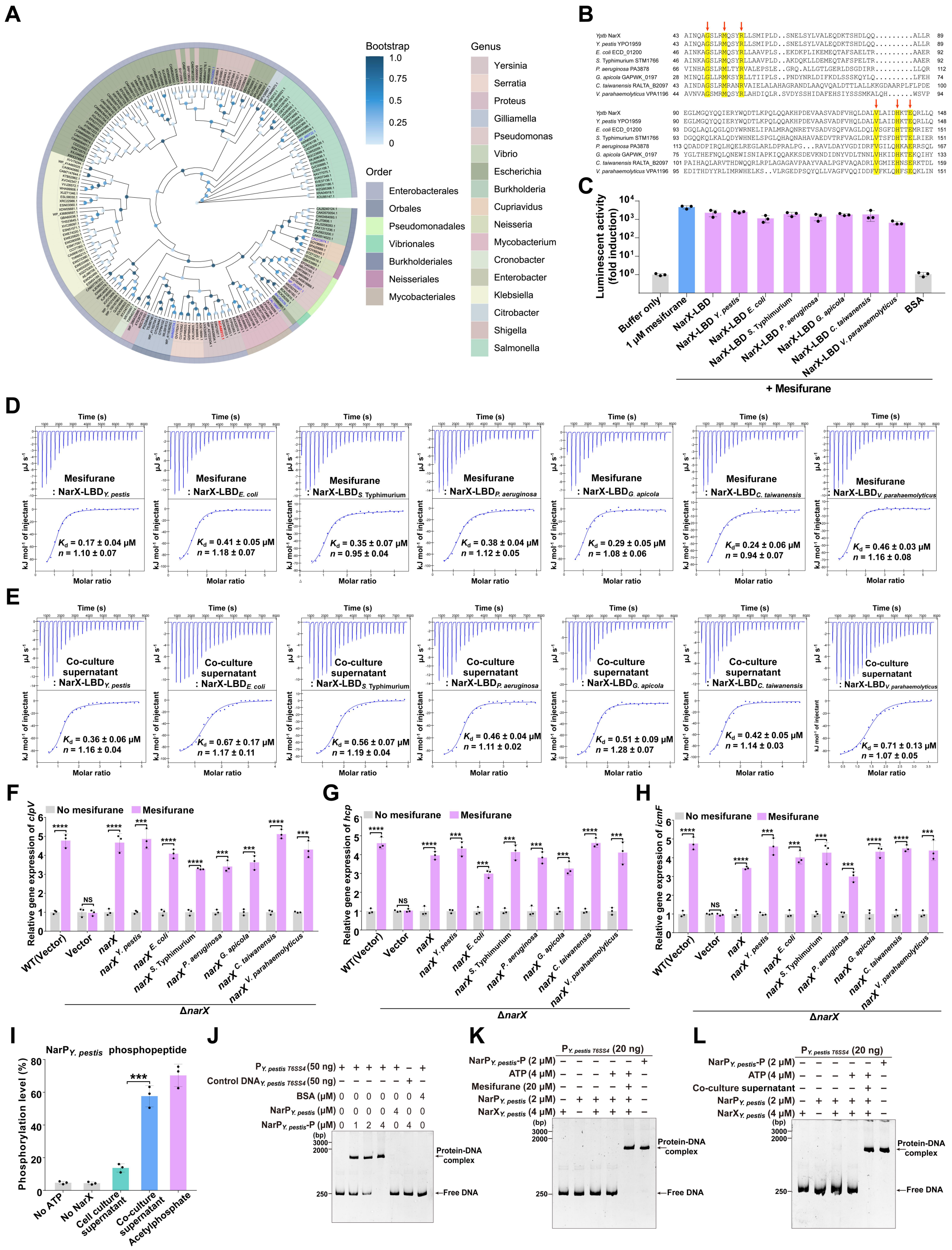

**Figure S5. Mesifurane sensing by NarX is a conserved mechanism in bacteria.**

**(A)** Phylogenetic analysis of NarX homologs. A maximum-likelihood tree was constructed from 200 NarX sequences. The *Yptb* NarX is highlighted in red, and experimentally verified homologs are in blue. Outer rings indicate bacterial taxonomy at the order level. Bootstrap values are visualized as a blue gradient.

**(B)** Sequence alignment of NarX ligand-binding domains. Amino acid sequences of NarX-LBD and homologous LBDs were aligned using ClustalW and visualized with GeneDoc 2.7. Highly conserved residues G48, M52, R56, V135, H140, and E143 are highlighted in yellow.

**(C)** NarX homologs bind the mesifurane. Purified NarX proteins from *Yersinia pestis* CO92, *Escherichia coli* BL21, *Salmonella* Typhimurium SL1344, *Pseudomonas aeruginosa* PAO1, *Gilliamella apicola* wkB1, *Cupriavidus taiwanensis* LMG19424 and *Vibrio parahaemolyticus* RIMD2210633 were incubated with mesifurane; captured ligand was heat-eluted and quantified via the *V. harveyi* MM32 bioluminescence assay. 1 μM mesifurane was a positive control, and BSA served as a negative control. Data are fold induction relative to the buffer control.

**(D and E)** ITC analysis of mesifurane (D)/co-culture supernatant (E) binding to NarX homologs. The displayed thermogram in (D, E) is one representative of three independent experiments; $K_d$ and binding stoichiometry (*n*) are presented as mean ± s.d. from three independent experiments.

**(F-H)** Heterologous *narX* genes restore mesifurane sensing in *Yptb* Δ*narX*. qRT-PCR analysis of T6SS gene expression for *clpV* (F), *hcp* (G), and *icmF* (H) in complemented *Yptb* Δ*narX* strains after stimulation with 10 μM mesifurane for 2 h. Expression was normalized to 16S rRNA and is presented as fold change relative to each unstimulated strain.

**(I)** Mesifurane stimulates NarP phosphorylation via *Y. pestis* NarX. Purified *Y. pestis* NarP and NarX were incubated with ATP in the presence or absence of co-culture supernatant for 40 min. Reactions containing acetyl phosphate served as a positive control for *Y. pestis* NarP phosphorylation; reactions lacking ATP or *Y. pestis* NarX served as a negative control. Phosphorylation of *Y. pestis* NarP was quantified by LC-MS/MS analysis.

**(J)** Phosphorylated *Y. pestis* NarP binds the T6SS-4 promoter. EMSA showing binding of acetyl phosphate-phosphorylated *Y. pestis* NarP to the *Y. pestis* T6SS-4 promoter probe. BSA and coding-region DNA fragments served as negative controls.

**(K and L)** *Y. pestis* NarP binds the T6SS-4 promoter in vitro. EMSA was performed with *Y. pestis* NarP and the T6SS-4 promoter probe in the presence or absence of mesifurane/co-culture supernatant and/or *Y. pestis* NarX. Acetyl phosphate-phosphorylated *Y. pestis* NarP was included as a positive control. The gel shown in (J-L) is representative of three independent experiments with similar results.

Data are represented as mean ± SD of three biological replicates, each with three technical replicates. Statistical significance was determined with the two-tailed unpaired Student's *t*-test. ***$P < 0.001$; ****$P < 0.0001$; NS, not significant.

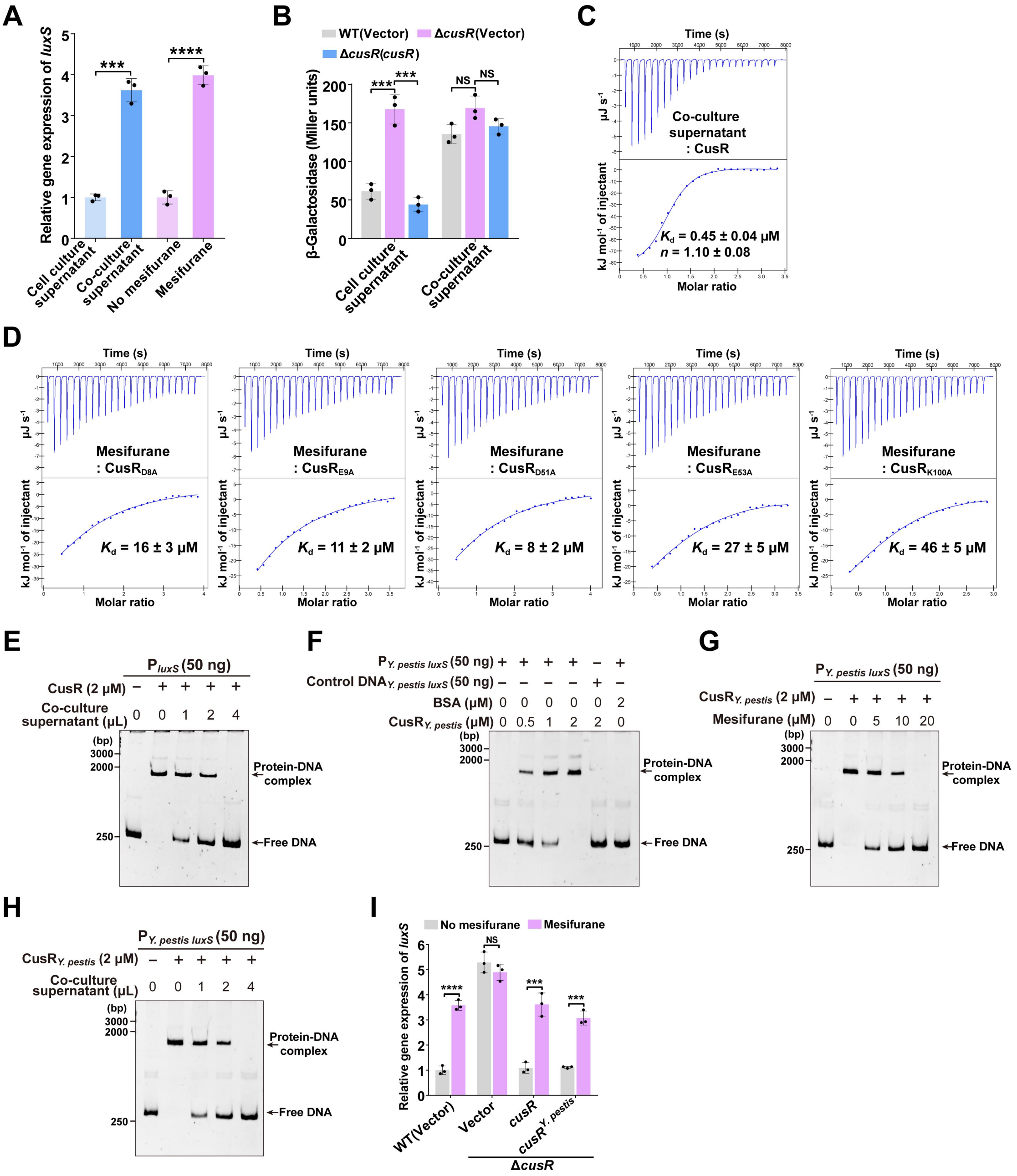


**Figure S6. CusR binding mesifurane signaling to derepress *luxS* expression.**

**(A)** qRT-PCR analysis of *luxS* genes in WT *Yptb* after 2 h of stimulation. Cells were treated with cell culture supernatant (unstimulated control), co-culture supernatant (containing mesifurane), or 10 μM mesifurane. Expression was normalized to 16S rRNA and is presented as fold change relative to the unstimulated control.

**(B)** β-Galactosidase assays quantifying $P_{T6SS4}$*::lacZ* reporter activity in the indicated *Yptb* strains treated with co-culture supernatant for 2 h. *Yptb* strains treated with cell culture supernatant served as a control.

**(C)** The binding affinity between CusR and co-culture supernatant was evaluated using ITC. The displayed thermogram is one representative of three independent experiments; $K_d$ and binding stoichiometry (*n*) are presented as mean ± s.d. from three independent experiments.

**(D)** ITC analysis of the binding affinity between CusR point mutants and mesifurane. The displayed thermogram is one representative of three independent experiments; $K_d$ and binding stoichiometry (*n*) are presented as mean ± s.d. from three independent experiments.

**(E)** EMSA for CusR binding to the *luxS* promoter in the presence or absence of co-culture supernatant. Co-culture supernatant was simultaneously added with CusR to the reaction system.

**(F)** EMSA for *Y. pestis* CusR binding to *Y. pestis luxS* promoter. BSA and DNA fragments amplified from the coding regions of the *Y. pestis luxS* gene were used as negative controls.

**(G and H)** EMSAs for *Y. pestis* CusR binding to the *Y. pestis luxS* promoter in the presence or absence of mesifurane (G) or co-culture supernatant (H). Co-culture supernatant or mesifurane was simultaneously added with CusR to the reaction system. The gels shown in (**E-H**) are representative of three independent experiments with similar results.

**(I)** *Y. pestis cusR* complements the *Yptb* Δ*cusR* mutant. qRT-PCR analysis of *luxS* expression in complemented strains after stimulation with 10 μM mesifurane for 2 h. Expression was normalized to 16S rRNA and is presented as fold change relative to the unstimulated WT control.

Data are represented as mean ±SD of three biological replicates, each with three technical replicates. Statistical significance was determined with the two-tailed unpaired Student's *t*-test. ****P* < 0.001; *****P* < 0.0001; NS, not significant.

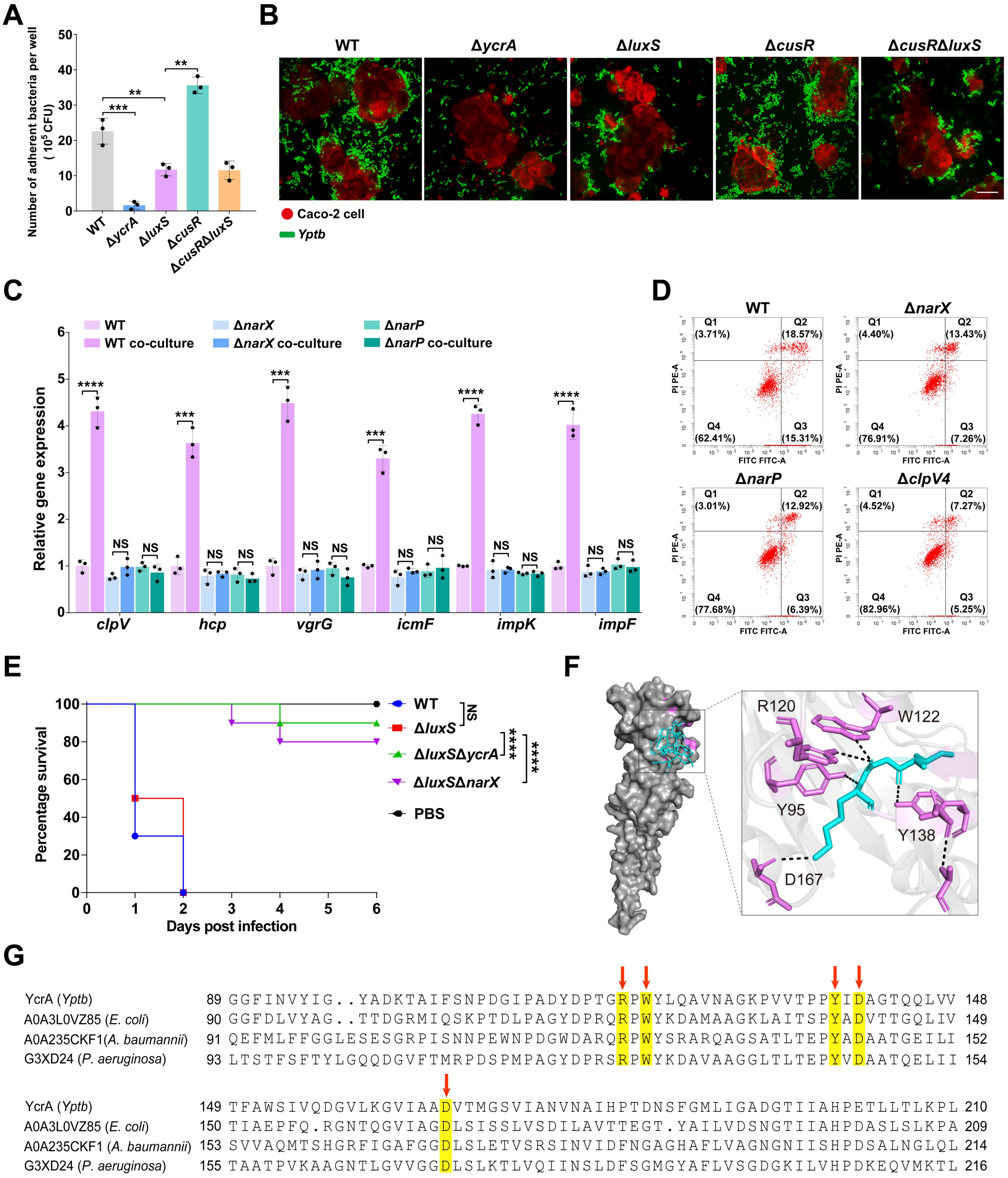


**Figure S7. "sense, arm, recruit" circuit is fully operational during infection of intestinal epithelial cells.**

**(A)** Quantification of *Yptb* adherence to intestinal epithelial cells. Caco-2 cells were infected with the indicated *Yptb* strains at MOI of 100 for 1 h. Nonadherent bacteria were removed by washing, and adherent bacteria were released by cell lysis, serially diluted, and enumerated by CFU plating.

**(B)** Caco-2 cells (CellTracker™ Deep Red, red) were infected with GFP-expressing *Yptb* strains (green) at MOI 100 for 1 h. Nonadherent bacteria were removed by washing prior to imaging. Scale bar, 20 μm. Images are representative of three independent experiments.

**(C)** T6SS gene expression during epithelial cell infection requires NarX and NarP. qRT-PCR analysis of T6SS gene expression in the indicated strains following infection of Caco-2 cells. Bacteria were added to macrophages at an MOI of 100, and after 2 h of co-culture, bacterial RNA was extracted for analysis. Expression was normalized to 16S rRNA and is presented as fold change relative to bacteria cultured without Caco-2 cells.

**(D)** Flow cytometric analysis of Caco-2 cell apoptosis. Caco-2 cells were infected with the indicated *Yptb* strains (MOI 100) for 2 h and stained with Annexin V-FITC/PI. A representative result from three independent experiments is shown.

**(E)** Survival curves of BALB/c mice following intraperitoneal infection with indicated *Yptb* strains ($1 \times 10^7$ CFU) (n = 10 mice per group).

**(F)** Predicted binding mode of peptide antagonist within the YcrA-LBD. Peptide antagonist is shown as cyan sticks and key interacting residues are shown as purple sticks, with dashed lines indicating potential hydrogen bonds.

**(G)** Sequence alignment of YcrA ligand-binding domains. Amino acid sequences from *Yptb*, carbapenem-resistant clinical isolates of *E. coli* (EC155), *A. baumannii* (Ab-C17), and *P. aeruginosa* (ATCC BAA-2108) were aligned using ClustalW and visualized with GeneDoc 2.7. Highly conserved residues R120, W122, Y138, D140, and D167 are highlighted in yellow.

For mouse survival assays (E), two independent experiments were performed; one representative experiment is shown, with the indicated number of mice per group. For (A and C), data are presented as mean ± SD of three biological replicates, each with three technical replicates. Statistical significance was determined with the two-tailed unpaired Student's *t*-test (A and C) or Log-rank (Mantel–Cox) test (E). ***$P < 0.001$; ****$P < 0.0001$; NS, not significant.

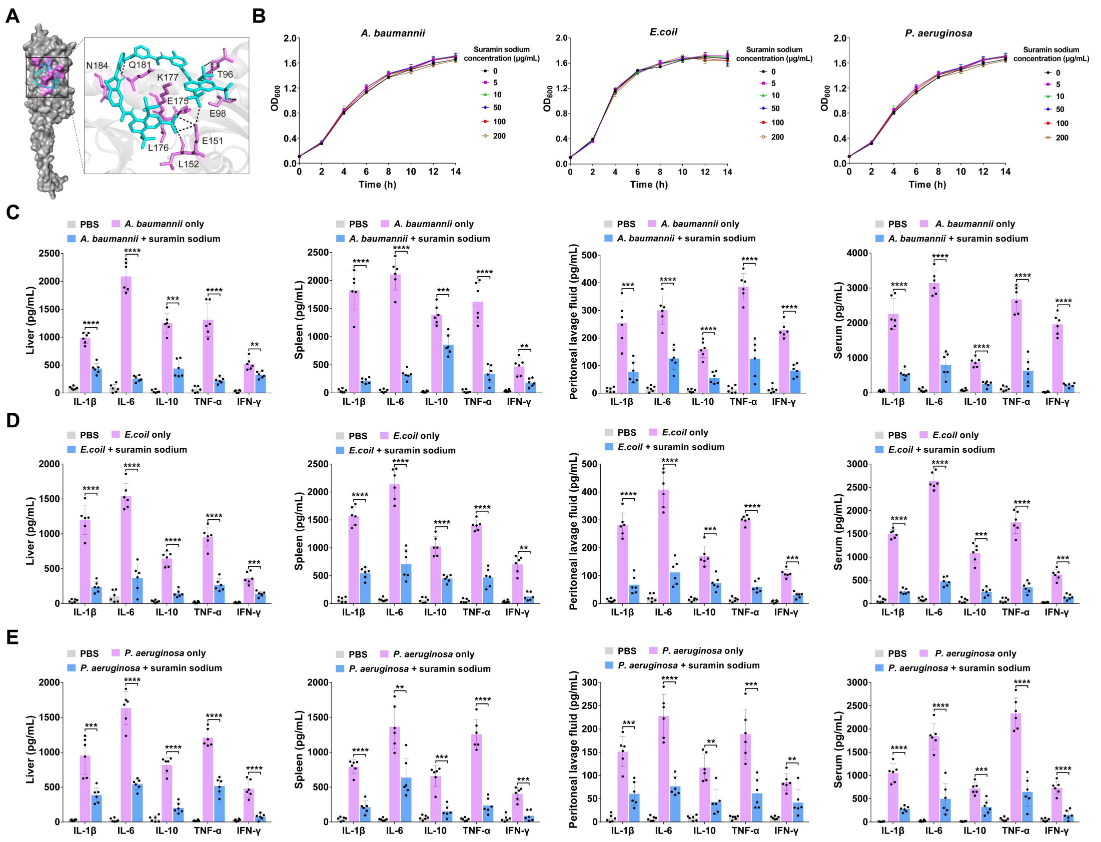


**Figure S8. Suramin sodium lacks direct antibacterial activity and suppresses infection-induced inflammation.**

**(A)** Predicted binding mode of suramin sodium within the YcrA-LBD of *P. aeruginosa*. Suramin sodium is shown as cyan sticks and key interacting residues are shown as purple sticks, with dashed lines indicating potential hydrogen bonds.

**(B)** Suramin sodium does not affect growth of clinical isolates. Growth curves of the indicated clinical isolates cultured in the presence of PBS or suramin sodium at the concentrations indicated.

**(C-E)** Cytokine levels in serum, peritoneal lavage fluid, spleen, and liver from mice infected with *A. baumannii* Ab-C17 ($1\times 10^6$ CFU) (C), *E. coli* EC155 ($2\times 10^7$ CFU) (D), or *P. aeruginosa* ATCC BAA-2108 ($5\times 10^6$ CFU) (E). Infected mice received suramin sodium (10 mg/kg) or PBS subcutaneously beginning 2 h post-infection, and tissues were collected 14 h post-infection. (n = 6 mice per group).

For ELISA assays (C–E), six biological replicates (individual mice) were analyzed, each assessed in three technical replicates. Data are shown as mean ± SD. Growth curves (B) data are presented as mean ± SD of three biological replicates, each with three technical replicates. Statistical significance was determined with the two-tailed unpaired Student's *t*-test. *$P < 0.1$; **$P < 0.01$; ***$P < 0.001$; ****$P < 0.0001$; NS, not significant.

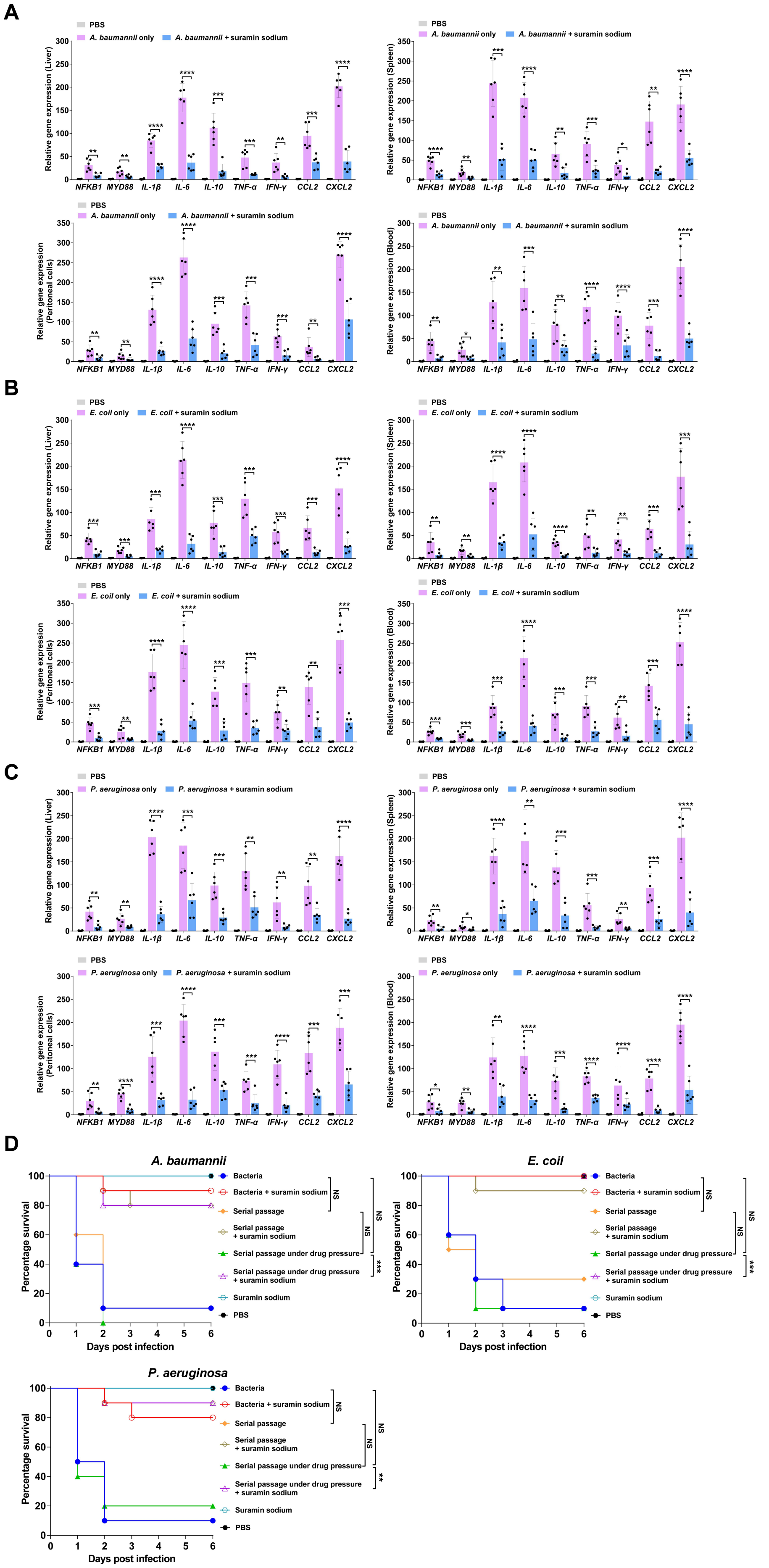

A
PBS
A. baumannii only
A. baumannii + suramin sodium
Relative gene expression (Liver)
Relative gene expression (Spleen)
Relative gene expression (Peritoneal cells)
Relative gene expression (Blood)
NFKB1
MYD88
IL-1β
IL-6
IL-10
TNF-α
IFN-γ
CCL2
CXCL2
B
E. coil only
E. coil + suramin sodium
C
P. aeruginosa only
P. aeruginosa + suramin sodium
D
A. baumannii
E. coil
P. aeruginosa
Percentage survival
Days post infection
Bacteria
Bacteria + suramin sodium
Serial passage
Serial passage + suramin sodium
Serial passage under drug pressure
Serial passage under drug pressure + suramin sodium
Suramin sodium
PBS
NS

**Figure S9. Suramin sodium suppresses proinflammatory transcription and does not select for resistance.**

**(A-C)** qRT-PCR analysis of proinflammatory and chemokine transcripts in spleen, liver, peritoneal cells, and blood from mice infected with *A. baumannii* Ab-C17 ($1\times 10^6$ CFU) (A), *E. coli* EC155 ($2\times 10^7$ CFU) (B), or *P. aeruginosa* ATCC BAA-2108 ($5\times 10^6$ CFU) (C). Infected mice received suramin sodium (10 mg/kg) or PBS subcutaneously beginning 2 h post-infection, and tissues were collected 14 h post-infection. (n = 6 mice per group). Expression was normalized to *Actb* and is presented as fold change relative to uninfected controls.

**(D)** Resistance development assay. Three clinical isolates were serially passaged for ~200 generations in the absence (serial passage) or presence (serial passage under drug pressure) of suramin sodium (10 μg/ml). The ancestral strain was maintained without passage. The ancestral strain and both passaged populations were then used to infect mice. Mice were infected with the indicated clinical isolates as in (A-C). Infected mice received suramin sodium (10 mg/kg) or PBS subcutaneously beginning 2 h post-infection, every 12 h for the first 3 days and every 24 h thereafter. Uninfected controls were given suramin sodium or PBS on the same schedule. Survival was monitored for 6 days. (n = 10 mice per group).

For mouse survival assays (D), two independent experiments were performed; one representative experiment is shown, with the indicated number of mice per group. For qRT-PCR assays (A–C), six biological replicates (individual mice) were analyzed, each assessed in three technical replicates. Data are presented as mean ± SD. Statistical significance was determined with the two-tailed unpaired Student's *t*-test (A–C) or Log-rank (Mantel–Cox) test (D). *$P < 0.1$; **$P < 0.01$; ***$P < 0.001$; ****$P < 0.0001$; NS, not significant.

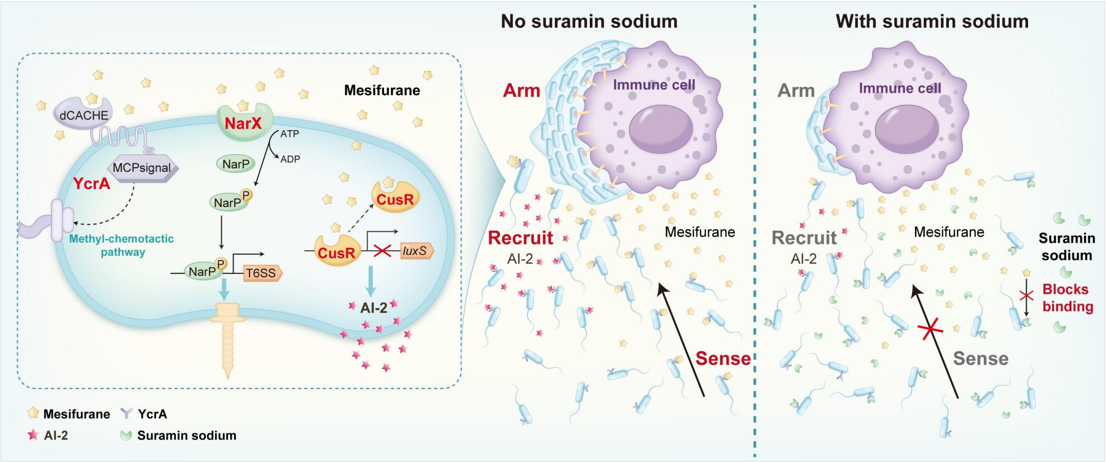


**Figure S10. Model of bacterial hijacking of a host metabolite to orchestrate infection, and its therapeutic disruption.**

*Yptb* intercepts the host-derived metabolite mesifurane through three distinct sensory systems that together execute a coordinated "Sense, Arm, Recruit" program. The chemoreceptor YcrA mediates chemotaxis toward host cells by detecting mesifurane, positioning the pathogen in proximity to its target. Upon arrival, the NarX-NarP two-component system perceives the host signal and licenses the expression and assembly of the T6SS, enabling contact-dependent killing. Concurrently, mesifurane antagonizes the transcriptional repressor CusR, relieving repression of *luxS* and amplifying endogenous AI-2 production. This AI-2 signal is secreted and recruits additional bacteria to the infection site, transforming a solitary encounter into a coordinated multi-bacterial attack. This tripartite circuit converges on a single druggable node: the central sensor YcrA. Suramin sodium, a repurposed drug, blocks the YcrA ligand-binding pocket, preventing mesifurane perception. Pharmacological blockade of YcrA with suramin sodium prevents mesifurane perception, disabling chemotaxis and thereby disrupting the coordinated attack program, without inhibiting bacterial growth. This work reveals how a single host metabolite can be co-opted to coordinate a multi-module offensive program, and demonstrates that disrupting its central sensor converts this circuitry into a therapeutic vulnerability—offering a path toward anti-infectives that target what a pathogen hears rather than whether it lives.
